# Genetic suppression of myeloid receptor Clec7a attenuates microglia neuroinflammation and promotes microglial phagocytosis to delay disease progression in ALS models

**DOI:** 10.64898/2026.05.04.722437

**Authors:** Xuan Chen, Huilan Yan, Haichao Wei, Sanaz Sajadi, Jingwen Hu, Vinicius Vasconcellos, Ashley Kim, Tarun Shriram, Huizhong Tan, Kyoeun Keum, Jiaqian Wu, Martin Paukert, Yongjie Yang

## Abstract

Microglial activation has been closely associated with accelerated ALS disease progression. However, specific microglial pathways that regulate microglial activation and ALS disease progression remain limitedly understood. Here, we determined the role of Clec7a (or Dectin-1), a core signature gene of disease-associated microglia (DAM) in ALS, in regulating microglial activation and ALS disease progression. Our spinal cord scRNA-Seq results found that Clec7a deficiency specifically attenuated microglial neuroimmune gene expression in SOD1G93A mice and human ALS. In addition, *in vivo* two-photon imaging of human (h) TDP43 phagocytosis by microglia in the cortex showed that Clec7a deficiency promotes microglial phagocytosis of pathological hTDP43 by enhancing microglial process dynamics. Subsequent survival analysis further showed that selective deletion of Clec7a in microglia mitigates motor neuron degeneration and delays disease progression in SOD1G93A ALS mice. Together, our results establish that Clec7a is a key regulator in shaping disease microglial functions and promotes disease progression in ALS.

## Introduction

Amyotrophic Lateral Sclerosis (ALS) is a fast-progressing and devastating motor neuron (MN) degenerative disease with limited treatment options. Pathological examinations and recent RNA sequencing of human ALS postmortem tissues^1^ and studies from ALS models have demonstrated that MN degeneration is caused by both cell-autonomous MN and non-cell autonomous glial mechanisms^2^. In particular, microglial activation in affected CNS regions, manifested by elevated neuroinflammation and impaired phagocytosis, has been consistently observed in human ALS and ALS models^3,4^, which is closely associated with accelerated ALS disease progression^1,5^. However, specific mechanisms that regulate microglial activation and microglial pathways that modulate ALS disease progression remain limitedly understood, except for the classical microglial NF-kB pathway^6^, which has been shown to induce microglial toxicity to MNs and contribute to ALS disease progression^6^. While ALS genes, such as *TBK1*, *OPTN*, and *C9ORF72*, have begun to implicate microglial pathways in ALS pathogenesis, RNA-seq^7,8^ and spatial transcriptome^9^ analysis of microglia in ALS and other neurodegenerative diseases^10^ have characterized distinct molecular signatures of disease-associated microglia (DAM), offering renewed insights in better understanding microglial mechanisms in ALS pathogenesis.

Clec7a (also named Dectin-1), a myeloid cell pattern recognition receptor^11^, has been identified as a core signature gene of DAM in neurodegenerative diseases including ALS^7,10^, and its upregulation in SOD1G93A ALS mice has been shown to depend on the TREM2-ApoE signaling^7^. Clec7a mRNA expression was found to be highly enriched in the cortex of human ALS-Glia subtype^1^ and also in ALS spinal cords which are inversely correlated with disease progression^5^. Interestingly, Clec7a expression is minimally detected in physiological microglia except transiently expressed in proliferative region-associated microglia (PAM) during development^12^. Although Clec7a is classically activated by a β-1,3-glucan structure (often found in fungal pathogens) in peripheral myeloid cells to induce cytokine release and to promote phagocytosis^13^, endogenous ligands for Clec7a in the CNS remain unknown, though N-glycosylated proteins that are highly present in the CNS pathology such as Galectin-9 and Vimentin have been proposed^14^. Clec7a has been shown to have context-dependent roles in neuropathology. Clec7a can be either neurotoxic or neuroprotective in CNS axon injury^14^, while Clec7a has been shown to limit autoimmune neuroinflammation in the EAE model^15^. On the other hand, microglial Clec7a mediates synapse elimination in ischemic stroke models to worsen neurobehavioral outcomes. In Alzheimer’s disease (AD), microglial Clec7a enhances microglial phagocytosis of Aβ plaques in the 5xFAD model^16,17^.

How selective Clec7a up-regulation in microglia impacts ALS pathogenesis remains little known. A recent study showed that Clec7a deficiency in MATR3 S85C ALS mice was found to have no impact on disease progression^18^; however, MATR3 S85C ALS mice are not representative of human ALS pathology in lack of MN loss in the motor cortex and spinal cord but instead have dramatic loss of Purkinje cells in cerebellum^19^. Given Clec7a’s context-dependent roles in CNS pathology and the close association between microglial activation and fast-progressive pathology in ALS^1^, it is essential to understand Clec7a’s roles in regulating microglia-mediated pathogenesis in ALS. In the current study, we investigated *in vivo* functions of disease-induced microglial Clec7a in regulating neuroinflammation, microglial phagocytosis of TDP43 aggregates, and ALS disease progression in SOD1G93A and TDP43 ALS models. Our results establish that Clec7a is a key regulator in shaping disease microglial functions and promotes disease progression in ALS.

## Results

### Disease-induced selective up-regulation of Clec7a protein in microglia in ALS mouse models and human ALS spinal cords

Although previous RNA-Seq analysis showed up-regulation of Clec7a mRNA in disease microglia in ALS^1,7,8^, whether Clec7a protein levels are increased in disease microglia in ALS models and especially in human ALS remained unexplored. Our Clec7a immunostaining found clear Clec7a expression (white arrows, Fig. 1a iv-vi) in ∼80% and 97% Iba1^+^ microglia at mid- (P115-120 old) and end-disease stages of SOD1G93A mice, respectively, but Clec7a was minimally detected in age-matched WT mice (yellow arrows, Fig. 1a i-iii). The percentage of Clec7a^+^ microglia (Clec7a^+^Iba1^+^) over total Iba1^+^ microglia was also gradually increased from pre-symptomatic (P70-80, ∼30%) to mid- (80%), and end-disease stages (∼P140, 97%) in spinal cords of SOD1G93A mice (Fig. 1b). In parallel, microglial Clec7a intensity (Fig. 1c) and total Clec7a levels in spinal cords (Extended Data Fig. 1a-b) were also increased as disease progresses. Induced Clec7a immunoreactivity was not found in GFAP^+^ astrocytes and Olig2^+^ oligodendrocytes (Extended Data Fig. 1c), nor in CD206^+^ border-associated macrophage (BAM, Extended Data Fig. 1d). Disease-induced selective up-regulation of Clec7a in microglia was also observed in the PFN1C71G ALS model (white arrows, Fig. 1d-e) and previously in the TDP-43 ΔNLS model^20^. To determine whether Clec7a protein is indeed increased in disease microglia in human ALS, we next performed Clec7a immunostaining on paraffin spinal cord sections of sporadic (s) and familial (f) ALS (both *SOD1* and *C9ORF72* mutations, Extended Data Fig. 1e) and age-matched healthy controls that were obtained from Target ALS consortium. Consistent with ALS mouse models, Clec7a positive labeling was frequently observed in both sALS (50%) and fALS (83%) (red arrows, Fig. 1f-g) with typical microglial morphology, but not in control sections (Fig. 1f-g).

**Figure 1.**
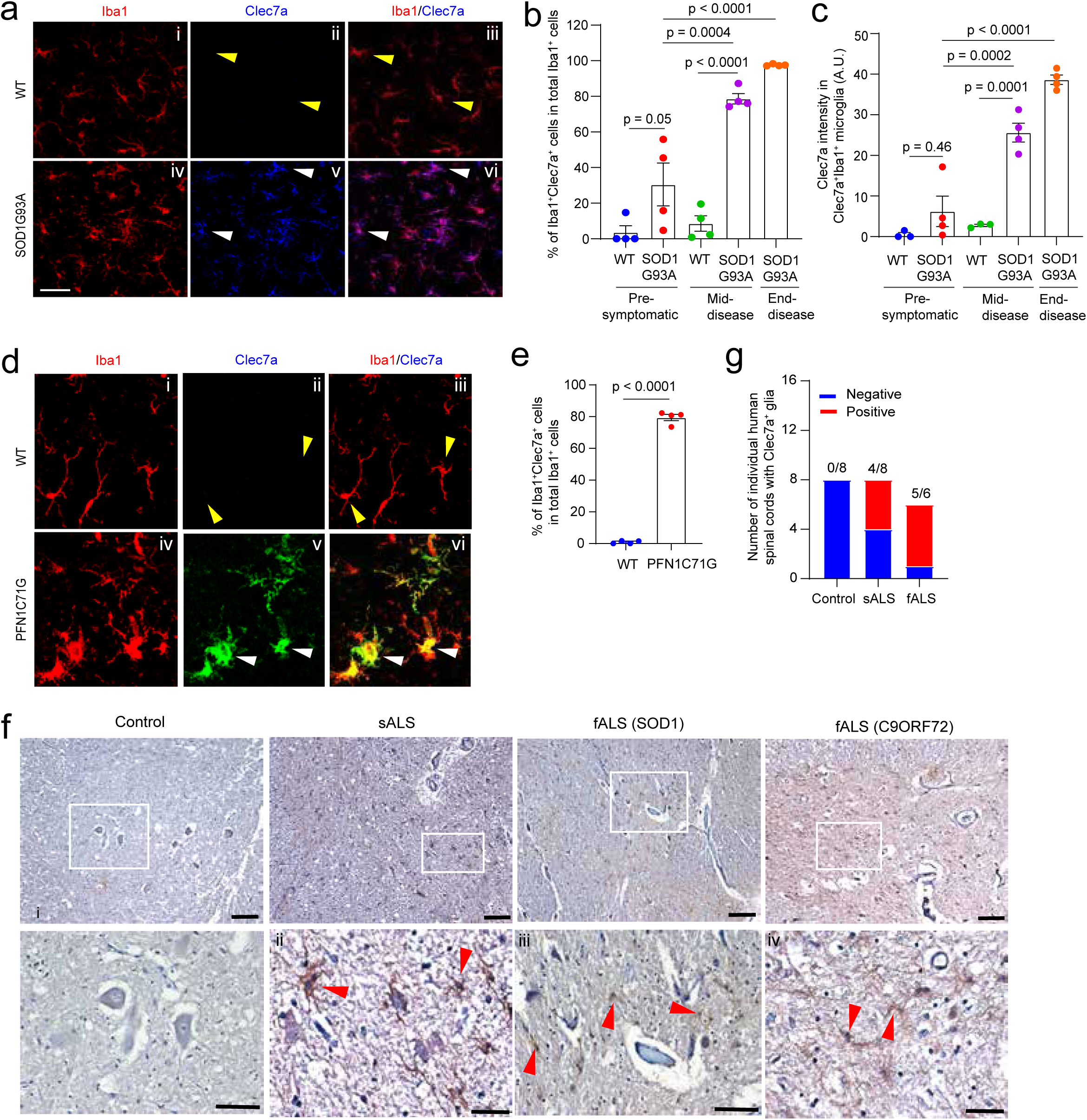
Selective up-regulation of Clec7a protein in spinal cord microglia of diseased SOD1G93A mice and human ALS. **a.** Representative images (40x) of Clec7a immunostaining in mid-stage (P110-115) SOD1G93A and age-matched control wild type (WT) mice. White arrows: Clec7a^+^Iba1^+^ microglia; Yellow arrows: Clec7a^-^Iba1^+^ microglia; Scale bar: 50μm; Quantification of the percentage (**b**) and Clec7a immunoreactivity (**c**) in Clec7a^+^Iba1^+^ microglia on spinal cord sections of SOD1G93A mice at pre-symptomatic (P70-80), mid- (P110-115), and end (∼P140)-stages. n = 6 images from 3-6 sections/mouse, 3-4 mice/group. p values were determined by one-way ANOVA and post-hoc Tukey’s analysis. Representative images (40x, **d**) of Clec7a and Iba1 immunostaining and quantification (**e**) of Iba1^+^Clec7a^+^ microglia in WT and PFN1C71G ALS mice. Scale bar: 20μm. n = 6 images/3 sections/mouse, 4 mice/group. p value determined by unpaired Student’s t-test. **f.** Representative Clec7a immunostaining DAB images of spinal cord sections from healthy control and ALS patients (sporadic, familial with SOD1 and C9ORF72 mutations). Green arrows: motor neurons; Red arrows: Clec7a^+^ glia cells; i-iv: magnified view of selected boxes from each upper panel image. Scale bar: 100μm (upper panel images) and 40 μm (i-iv). **g.** Quantification of the number of human control and ALS samples with Clec7a^+^ glia. Total 8 control, 8 sporadic (s) ALS, and 6 familial (f) ALS.

### Clec7a regulates microglial neuroimmune gene expression in spinal cords of diseased SOD1G93A mice and human ALS

Previous studies in peripheral immune cells have shown that spleen associated tyrosine kinase (SYK) is often activated downstream of Clec7a through its C-terminal immunoreceptor tyrosine-based activation motif (ITAM)^11^. SYK activation then signals through diverse downstream transcription pathways to trigger antifungal immune responses that involve reactive oxygen species (ROS) production, phagocytosis, and cytokine expressions^11^. We examined SYK activation by performing phosphorylated Y352 (p-Y352) SYK immunostaining, which has been shown to indicate early SYK activation^21^. We first confirmed that p-Y352 SYK is indeed able to detect activated SYK in LPS-stimulated primary microglia (Extended Data Fig. 2a-b), as previously described^22^. Specific p-Y352 SYK was observed in Iba1^+^ microglia across all experimental groups (Extended Data Fig. 2c). However, while Clec7a expression is increased in SDO1G93A mice as disease progresses as shown in Fig. 1, we observed no changes of the p-Y352 SYK intensity nor the number of p-352 SYK^+^Iba1^+^ microglia in all groups including in SOD1G93A^+^Clec7a^-/-^ mice (Extended data Fig. 2d-e), suggesting that deletion of Clec7a in SOD1G93A mice had no effect of altering p-Y352 SYK levels and downstream SYK activation was unlikely involved in Clec7a^+^ microglial functions in ALS SOD1 mice. To examine the effect of Clec7a on microglia neuroimmune signaling in ALS, we performed single-cell RNA sequencing (scRNA-Seq) analysis from spinal cords of WT, Clec7a^-/-^, SOD1G93A^+^, and SOD1G93A^+^Clec7a^-/-^ mice at mid-disease (P115-118 old) using the 10x genomics and Illumina pipeline (Extended Data Fig. 3a). Clec7a^-/-^ mice behave and breed normally with unchanged splenocytes except for impaired responses to fungal infection^13^. A total of 95,368 cells (3-4 mice/group) were sequenced and analyzed (Extended data Fig. 3b). The uniform manifold approximation and projection (UMAP) graphs from 4 groups all showed clear separation of major CNS cell types (Fig. 2a) based on representative genes (stacked violin plot in Extended Data Fig. 3c). The specificity of the data was confirmed by high Clec7a levels only in SOD1G93A^+^ mice but not in WT (minimal Clec7a levels), Clec7a^-/-^, and SOD1G93A^+^Clec7a^-/-^ mice, while high SOD1 levels (endogenous mouse and transgenic human SOD1 quantified together) were observed in SOD1G93A^+^ and SOD1G93A^+^Clec7a^-/-^ mice (Extended Data Fig. 3d), as expected. A clearly increased microglia population was observed in both SOD1G93A^+^ and SOD1G93A^+^Clec7a^-/-^ mice (Fig. 2b), suggesting disease-induced active microglial proliferation^4^.

**Figure 2.**
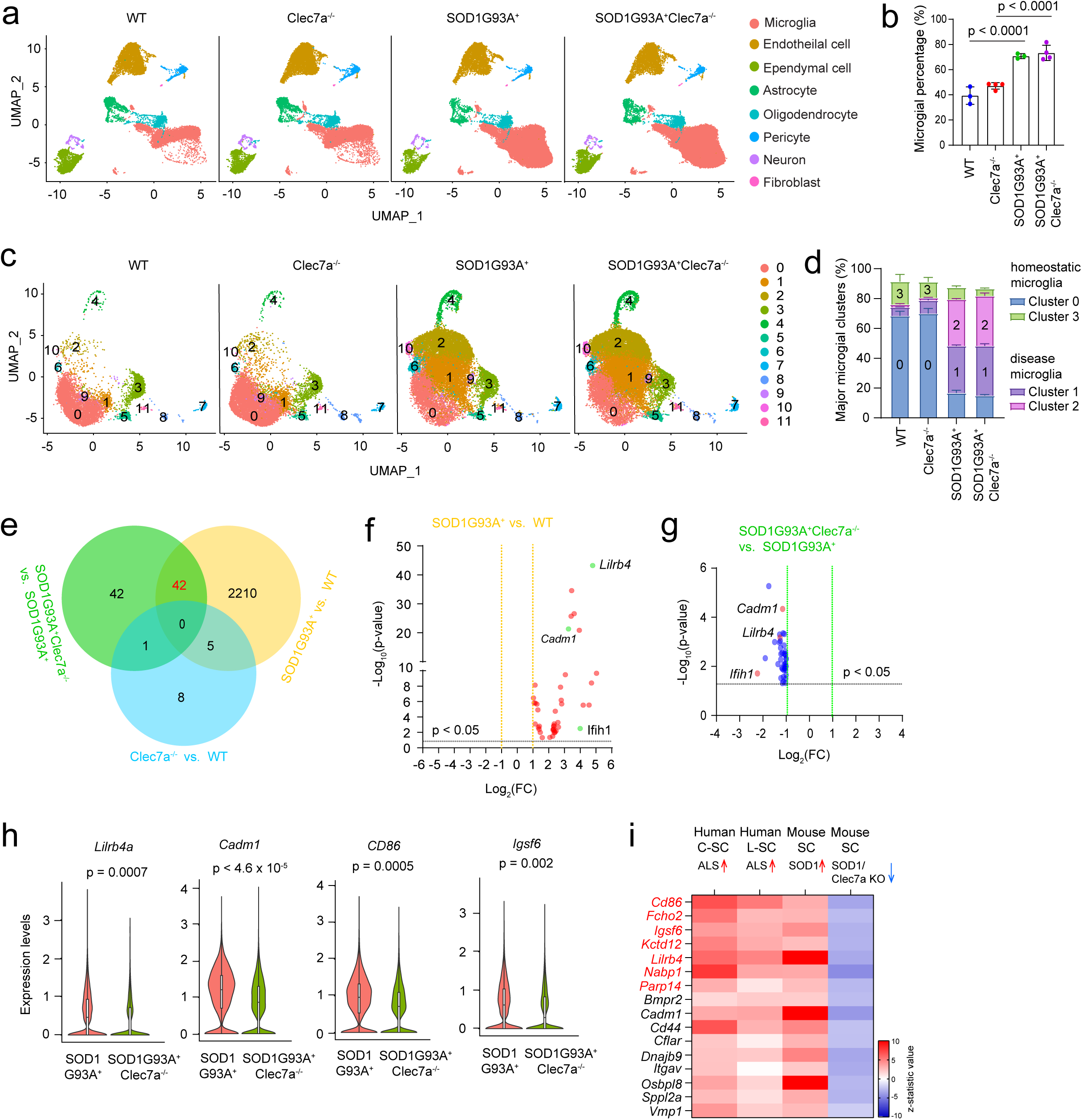
Single-cell RNA sequencing (scRNA-seq) identified Clec7a-regulated microglia neuroimmune genes in spinal cords of diseased SOD1G93A mice and human ALS. **a.** UMAP plot of major CNS cell types from each experimental group based on scRNA-seq data. **b.** Percentage of microglia in total analyzed cells from scRNA-seq in each experimental group. n = 3-4 mice/group. p values were determined by one-way ANOVA and post-hoc Tukey’s analysis. Clec7a KO mice are normal without obvious adverse phenotypes including motor deficits, as previously described^13^ and also confirmed by our own observations. UMAP of microglia subclusters (**c**, 12 identified) and the percentage changes (**d**) of major microglial subclusters (0, 1, 2, and 3) from all experimental groups. 0+3: homeostatic subclusters; 1+2: disease subclusters. n = 3-4 mice/group. **e.** Three-way Venn diagrams of SOD1G93A^+^Clec7a^-/-^ vs. SOD1G93A^+^, WT vs. SOD1G93A^+^, and WT vs. Clec7a^-/-^ comparisons. DEGs in each pair comparison were determined based on these criteria: p < 0.05, fold change (FC) > 2 or < -2, log_e_ (mean gene reads per sample) > 0.5. All 42 identified Clec7a-M-DEGs were up-regulated in SOD1G93A^+^ relative to WT (**f**) but down-regulated in SOD1G93A^+^Clec7a^-/-^ relative to SOD1G93A^+^ mice (**g**). Representative Clec7a-M-DEGs were highlighted. **h.** Expression level violin plots of representative Clec7a-M-DEG neuroimmune genes from scRNA-seq data of SOD1G93A^+^Clec7a^-/-^ and SOD1G93A^+^ mice. **i.** Overlap between Clec7a-M-DEGs and up-regulated neuroimmune genes in human ALS. SC: spinal cord; red arrow: up-regulated genes; blue arrow: down-regulated genes; genes highlighted in red are also reversely correlated with ALS disease duration based on^5^.

We next identified 12 distinct subclusters of microglia-like cells in experimental groups (Fig. 2c, also in Extended data Fig. 3e). Gene set enrichment analysis (GSEA) was performed to characterize their top functional pathways from which homeostatic microglia (primarily subclusters 0 and 3) and DAM (primarily subclusters 1 and 2) populations were identified. As expected, homeostatic microglia were significantly transitioned to disease microglia (Fig. 2d) in SOD1G93A^+^ and SOD1G93A^+^Clec7a^-/-^ mice. On the other hand, only an average 1.2% of peripheral macrophages (subcluster 7) and BAM (subcluster 8) were found based on specific gene transcripts (Extended data Fig. 3e) with no changes across groups (Fig. 2c). Other subclusters are likely to be in different transition states from homeostatic to disease microglia. As we observed no overall subcluster changes among groups, we decided to first obtain a full view of the overall microglial differentially expressed genes (DEGs) between SOD1G93A^+^ vs. SOD1G93A^+^Clec7a^-/-^, WT vs. SOD1G93A^+^, and WT vs. Clec7a^-/-^, using pseudo bulk analysis. Consistent with previous studies^8^, disease induced massive transcriptional changes (2257 DEGs, Supplementary Table 1) in microglia between WT and SOD1G93A (Extended Data Fig. 3f). Meanwhile, 85 DEGs, all down-regulated, were found in SOD1G93A^+^Clec7a^-/-^ microglia compared to SOD1G93A^+^ microglia (Extended Data Fig. 3g, Supplementary Table 2). While expression of *Trem2* and *Apoe* was significantly increased from WT to SOD1G93A mice, as previously reported^7,8^, their expression levels were unchanged from SOD1G93A to SOD1G93A^+^Clec7a^-/-^ mice (Extended Data Fig. 3h). Interestingly, these 85 significant DEGs were functionally clustered in several neuroimmune signaling pathways based on Ingenuity pathway analysis (Extended Data Fig. 3i). We also generated three-way Venn diagrams to identify overlapped microglial DEGs in these three comparisons (Fig. 2e) and found most overlapped DEGs between SOD1G93A^+^ vs. SOD1G93A^+^Clec7a^-/-^ and WT vs. SOD1G93A^+^ (Fig. 2e). Strikingly, all 42 overlapped microglial DEGs that were up-regulated in SOD1G93A^+^ relative to WT mice (Fig. 2f) were down-regulated in SOD1G93A^+^Clec7a^-/-^ relative to SOD1G93A^+^ mice (Fig. 2g), suggesting that they were disease-induced and were specifically reduced by the loss of Clec7a in disease microglia. We therefore named them Clec7a regulated microglial differentially expressed genes (Clec7a-M-DEGs, Supplementary table 3), which include neuroimmune genes (*Ifih1*, *Cd44*, *Cd86*, *Parp14*, *Igsf6*, *Itgav*, and *Sppl2a, etc.*), previously identified DAM genes (*Cadm1, Lilrb4*, also neuroimmune genes)^7^, and transcription factors (*Zeb2, Zswim6, and Zfp62*). Meanwhile, Clec7a-M-DEGs were not overlapping with DEGs identified between WT vs. Clec7a^-/-^ (only 14 total, Supplementary Table 4), suggesting that their downregulation in SOD1G93A^+^Clec7a^-/-^ relative to SOD1G93A^+^ mice is specifically due to the loss of disease-induced Clec7a. Expression changes of representative Clec7a-M-DEGs in SOD1G93A^+^ vs. SOD1G93A^+^Clec7a^-/-^ mice were shown in Fig. 2h . Our identified Clec7a-M-DEGs were also largely (25/42, 59%) detected in human spinal cords and majority (16/25, 64%) of detected Clec7a-M-DEGs were also significantly up-regulated in human ALS spinal cords^5^ (Fig. 2i), from which 7 neuroimmune genes (highlighted in red, Fig. 2i) were further reversely correlated with ALS disease duration^5^. These results showed that Clec7a-M-DEGs may directly contribute to ALS disease progression and loss of disease-induced Clec7a significantly decreases expression of microglial neuroimmune genes induced not only in SOD1G93A ALS mice but also in human ALS, thus disease-induced Clec7a may act as an upstream signal to promote microglia-mediated neuroimmune response in ALS.

To begin analyzing downstream transcriptional regulation of Clec7a-M-DEGs, we searched the established chromatin immunoprecipitation (ChIP) database TFLink^23^ and found that promoters of 50% (21/42, Supplementary Table 5) Clec7a-M-DEG can be bound by the microglial enriched transcription factor Mef2a (Fig. 3a). Interestingly, *Mef2a* is particularly induced in disease microglia (subclusters 1 and 2 in Fig. 2c) in SOD1G93A^+^ from WT mice (red dashed circle, Fig. 3b), while its expression in microglia was significantly (p = 0.03) down-regulated (1.8 fold, Fig. 3a, also see violin plot in Extended Data Fig. 3j) in SOD1G93A^+^Clec7a^-/-^ relative to SOD1G93A^+^ mice. This appears to be specific to *Mef2a*, as other transcription factors, such as *NF-kB*, that is classically downstream of Clec7a activation^11^ and has been shown to regulate microglial activation in ALS^6^, was not significantly (p = 0.26) changed and is only predicted to bind to promoters of 9 (out of 42) Clec7a-M-DEGs. In addition, the transcription factor *Zeb2* within Clec7a-M-DEGs and another similarly changed transcriptional factor *Sall 1*, important for microglial identity^24^, only potentially bind < 3 of Clec7a-M-DEG promoters (Fig. 3a). Subsequent Mef2a immunostaining confirmed its reduced expression in spinal cord microglia (white arrows, Fig. 3c) of SOD1G93A^+^Clec7a^-/-^ relative to SOD1G93A^+^ mice (quantification in Fig. 3d). As microglia are the primary CNS cell type to mediate elevated neuroinflammation and undergo microgliosis in ALS, we next examined microgliosis based on Iba1 immunostaining (representative images in Fig. 3e) in spinal cords of mice from experimental groups. As expected, Iba1^+^ microglial numbers were highly increased from WT to SOD1G93A^+^ mice but were significantly reduced (Fig. 3f) in SOD1G93A^+^Clec7a^-/-^ relative to SOD1G93A^+^ mice specifically at the mid-disease (P115) but not the pre-symptomatic (P80) stage (Fig. 3g). We also determined levels of pro-inflammatory cytokines such as IL-1β and TNFα that were considered to be primarily released by activated microglia^25^ and previously found to be well increased in spinal cords of SOD1G93A mice^8^ and human ALS^3^. Indeed, *IL-1β* and *TNF-α* mRNA levels were both increased in mid- (P115) and late-disease (P130) in SOD1G93A mice compared to control mice (Fig. 3h-i); however, their expression levels were significantly reduced in SOD1G93A^+^Clec7a^-/-^ mice at the late-disease stage (Fig. 3h-i).

**Figure 3.**
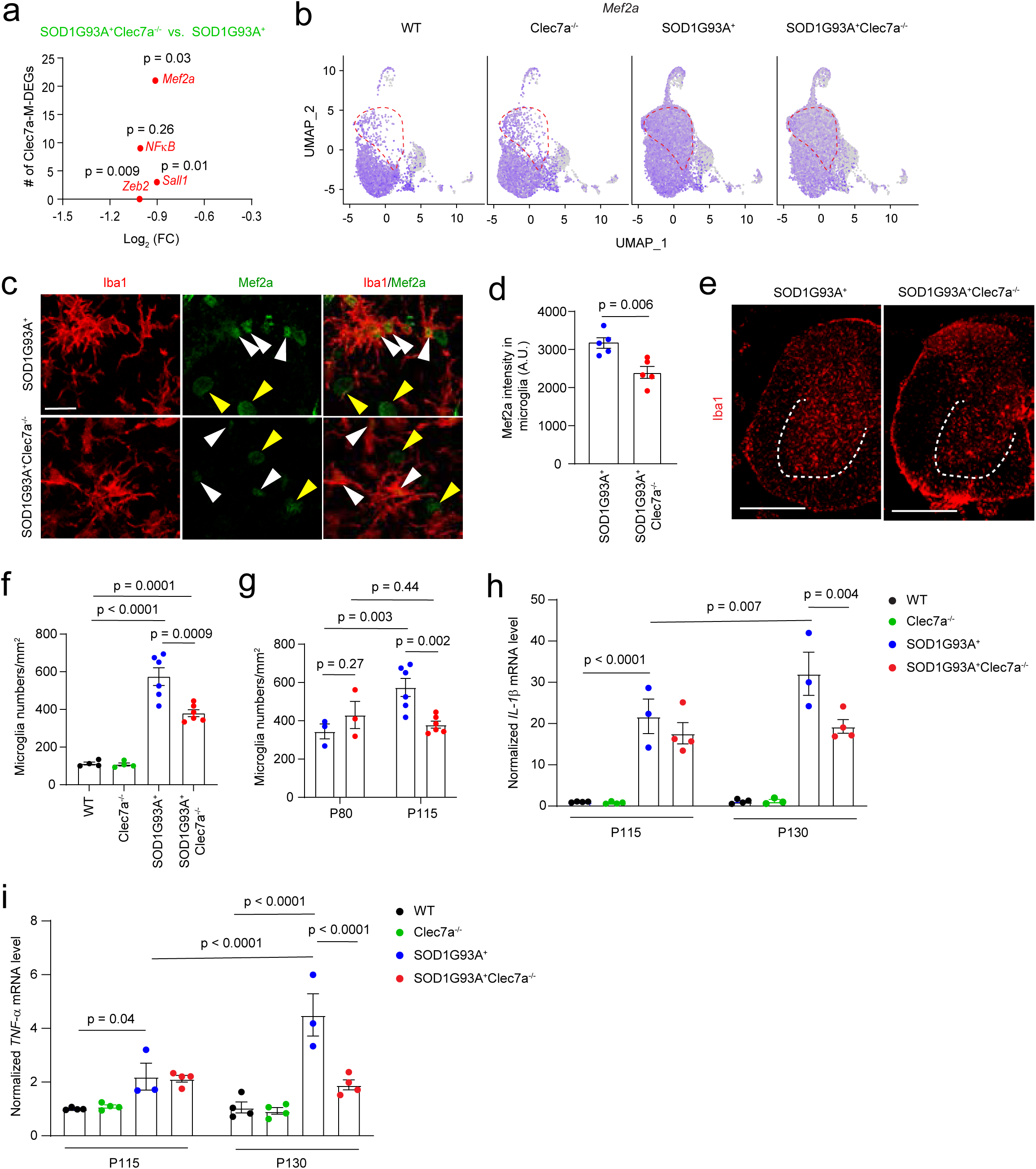
Clec7a deficiency suppresses microgliosis and neuroinflammation in spinal cords of diseased SOD1G93A mice. **a.** Number of genes in Clec7a-M-DEGs whose promoter sequence predicted to bind to selected transcription factors. The x axis showed reduced fold changes (Log_2_(FC)) of each transcription factor in SOD1G93A^+^Clec7a^-/-^ relative to SOD1G93A^+^ mice. The corresponding p values were shown next to each transcription factor. **b.** UMAP plot of *Mef2a* expression in microglial subclusters of each experimental group. Representative images (63x, **c**) and quantification (**d**) of Mef2a immunostaining in spinal cords of mid-stage (P115) SOD1G93A^+^Clec7a^-/-^ and SOD1G93A^+^ mice. White arrows: Mef2a immunoreactivity in microglia; Yellow arrows: Mef2a immunoreactivity in non-microglial cells; Scale bar: 20μm. n = 6 images from 3-6 sections/mouse, 5 mice/group. p values were determined by unpaired Student’s t-test. **e.** Representative images (40x) of Iba1 immunostaining in spinal cords of mid-stage (P115) SOD1G93A^+^Clec7a^-/-^ and SOD1G93A^+^ mice. Quantification of Iba1^+^ microglial numbers in all experimental groups (**f**) and during disease progression (**g**). n = 6 images from 3-6 sections/mouse, 4-6 mice/group. p values were determined by one-way ANOVA and post-hoc Tukey’s analysis. mRNA expression levels of pro-inflammatory cytokines *IL-1β* (**h**) and *TNF-α* (**i**) in all experimental groups during disease progression. n = 3-4 mice/group. p values were determined by one-way ANOVA and post-hoc Tukey’s analysis.

In contrast to the dramatic changes of microglial gene expression in SOD1G93A^+^Clec7a^-/-^ relative to SOD1G93A^+^ mice, astroglial genes were minimally impacted by the loss of Clec7a in SOD1G93A^+^ mice. Astroglial subclusters remained little changed among different groups (Extended Data Fig. 4a). GSEA analysis further characterized astrocyte subclusters and dramatic shift of resting/functional astrocytes (subclusters 0, 5, 6, and 7) to reactive/inflammatory astrocytes (subclusters 1 and 2) between WT and SOD1G93A mice, regardless of Clec7a genotype, was observed (Extended Data Fig. 4b). Only 2 DEGs (*Cx3cl10* and *Ccl2*) were found to be reduced in SOD1G93A^+^Clec7a^-/-^ relative to SOD1G93A^+^ mice while large number of DEGs were found between WT and SOD1G93A mice (Extended Data Fig. 4c, Supplementary Tables 6-8). Similarly, astrogliosis remained unchanged in spinal cords of SOD1G93A^+^Clec7a^-/-^ compared to SOD1G93A^+^ mice (Extended Data Fig. 4d-e). Overall, our results consistently suggest that deletion of disease-induced Clec7a ameliorates microgliosis and attenuates microglia-mediated neuroinflammation in spinal cords of SOD1G93A mice.

### Clec7a deficiency promotes microglial phagocytosis of pathological hTDP43 by enhancing microglial process dynamics in the motor cortex

Microglial phagocytosis is a critical clearance mechanism in CNS development and pathology to remove pathogens, degenerated neurons, myelin, and abnormal protein aggregates^26^. Several ALS associated (mutant) proteins, including SOD1, FUS, TDP43, C9ORF72, have been shown to form abnormal protein aggregates from which some further spread as disease progresses^27,28^. Whether Clec7a regulates microglial phagocytosis of ALS-relevant protein aggregates remains unexplored. As mutant SOD1 proteins are ubiquitously expressed in many cell types including microglia in SOD1G93A mice, it becomes unfeasible to examine microglial phagocytosis in SOD1G93A^+^Clec7a^-/-^ mice. On the other hand, TDP43 aggregates have been widely observed in human ALS pathology and are pathogenic in ALS^29^. We then decided to examine whether Clec7a deficiency affects phagocytosis of TDP43 aggregates. we obtained AAV9-hTDP43-GFP virus^30^ that allows focal overexpression of GFP-tagged human (h)TDP43 following the stereotaxic injection in the motor cortex (Fig. 4a). Overexpression of WT TDP43 has been previously shown to form aggregates^31^. We performed injections into the motor cortex but not into spinal cords in order to minimize tissue damage during surgery which can significantly complicate data interpretation. AAV9-hTDP43-GFP-transduced areas and expression levels of hTDP43-GFP are similar in both WT and Clec7a KO mice (Extended Data Fig. 5a). Expression of hTDP43 was mostly observed in cortical layers II-IV (within the white oval circle, Fig. 4ai) and was primarily overlapped with NeuN^+^ cells (magnified view, Fig. 4aii, ∼70% from quantification) with a predominant nuclear localization (70-80%, Extended Data Fig. 5b). About 5-7% of hTDP43-GFP was also found in Olig2^+^ oligodendrocytes but not in GFAP^+^ astrocytes (Extended Data Fig. 5c). Expressed hTDP43-GFP is also well-overlapped with hTDP43 immunoreactivity (Extended Data Fig. 5d), confirming the hTDP43 expression. The expression of hTDP43-GFP, but not control GFP, selectively and significantly induces Clec7a expression in microglia (white arrows, Fig. 4b, quantification in Fig. 4c, individual channel in Extended Data Fig. 5e). In parallel with Clec7a up-regulation, microglia around hTDP43-GFP were also more activated based on increased Iba1 immunoreactivity compared to GFP expression alone (Extended Data Fig. 5f). Consistently, hTDP43-GFP, but not control GFP, also induced a significant increase of p62 immunoreactivity (Extended Data Fig. 5g-h), an indication of abnormal protein aggregate formation and neuronal stress^32^. In parallel, immunostaining of phosphorylated S409/S410 TDP-43, a highly specific and consistent indicator for pathological, aggregated TDP-43 inclusions^33^, was also found to overlap well with hTDP43-GFP (Fig. 4di and enlarged boxes a and b) and could be found engulfed by Iba1^+^ microglia (Fig. 4d, enlarged box ai-avi). These results suggest that overexpressed hTDP43-GFP induces hTDP43 aggregates that trigger neuronal stress.

**Figure 4.**
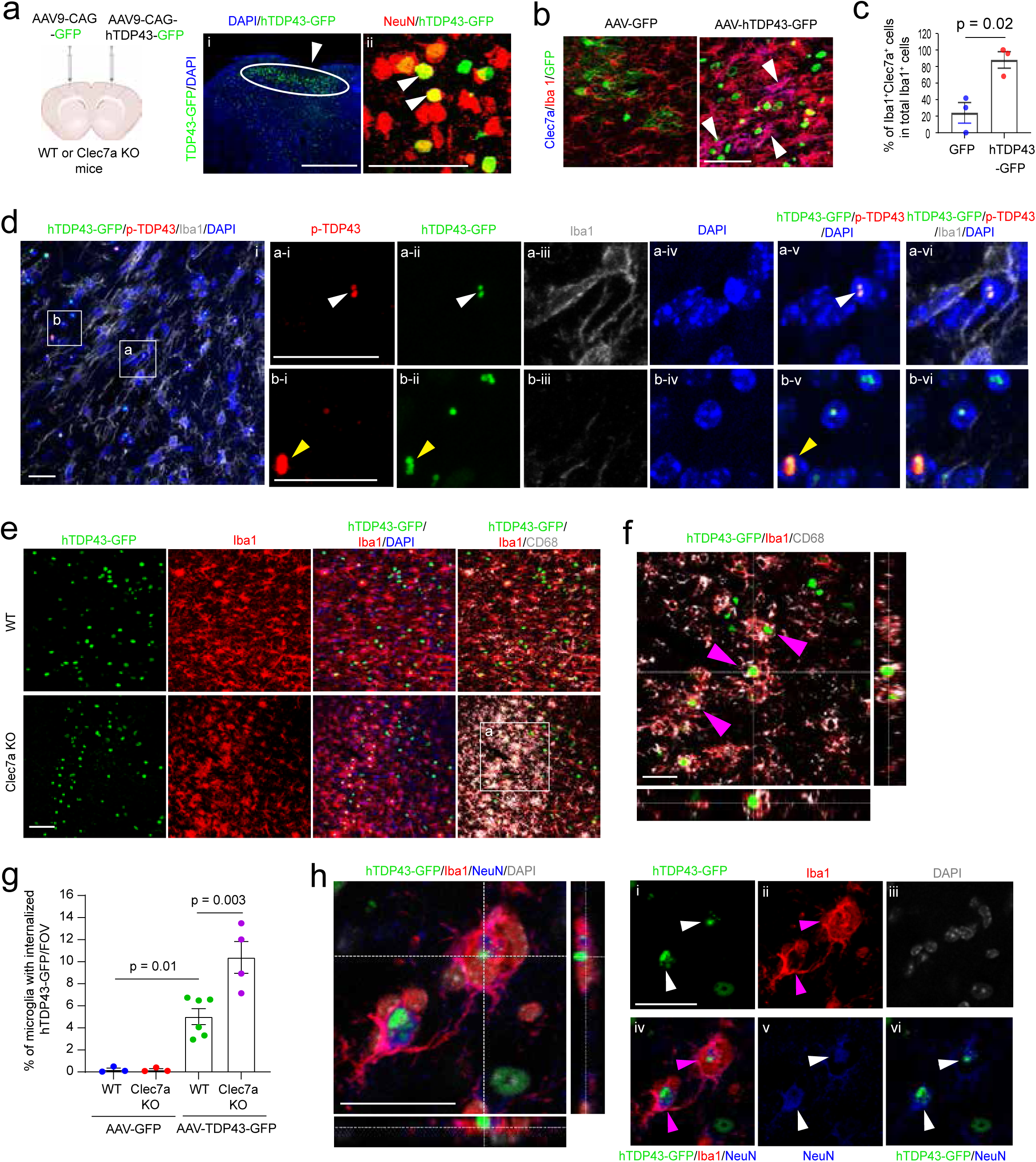
Clec7a deficiency promotes microglial phagocytosis of phosphorylated (S409/S410) and pathological hTDP43 in the motor cortex. **a.** Injection paradigm and representative images (20x in i, 40x in ii) to show a focal injection (white arrow in i) of AAV9-hTDP43-GFP into the motor cortex of WT mice (P90). The magnified view (ii) from i shows the co-localization of hTDP43-GFP with NeuN (white arrows). Scale bar: 100μm (i) and 50 μm (ii). Representative images (40x, **b**) of Clec7a immunostaining and quantification (**c**) of the percentage of Clec7a^+^Iba1^+^ microglia over total Iba1^+^ microglia on the motor cortex sections of control AAV9-GFP and AAV9-hTDP43-GFP-injected WT mice. White arrows: Clec7a^+^Iba1^+^ microglia; Scale bar: 50μm. n = 6-10 images from 3-6 sections/mouse, 3 mice/group, p values were determined by unpaired Student’s t-test. **d.** Representative images (63x) of phosphorylated (S409/S410) TDP43 immunoreactivity that overlaps with nuclear hTDP43-GFP and can be surrounded by microglia. box a: example of nuclear overlapped p-TDP43 and hTDP43-GFP (white arrows) engulfed by microglia; box b: example of non-nuclear overlapped p-TDP43 and hTDP43-GFP (yellow arrows) without microglia nearby; ai-avi and bi-bvi: individual and overlapped channels for p-TDP43, hTDP43, Iba1, and DAPI. Scale bar: 20μm (i) and 20μm (ai-avi and bi-bvi). **e.** Representative images (40x) of hTDP43-GFP phagocytosis by microglia in the motor cortex of WT and Clec7a KO mice. Scale bar: 50μm; **f.** Representative orthogonal view of the single focal plane (from the box a in Fig. 4e) showing internalized hTDP43-GFP inside CD68^+^Iba1^+^ microglia (indicated by purple arrows). Scale bar: 20μm. **g.** The percentage of microglia with internalized hTDP43-GFP per image in the motor cortex of WT and Clec7a KO mice. Images were taken from proximity (300-400μm) of the injected area but avoided the injection track. n = 6-12 images from 4-6 sections/mouse, 3-6 mice/group. p values were determined by one-way ANOVA and post-hoc Tukey’s analysis. **h,** Representative orthogonal view (40x) of the single focal plane showing hTDP43^+^NeuN^+^ neurons phagocytosed by microglia in the motor cortex of Clec7a KO mice. i-vi: individual and overlapped channels of hTDP43-GFP, Iba1, NeuN, and DAPI. Scale bar: 25μm (orthogonal view and i-vi).

By examining the co-localization of hTDP43-GFP with Iba1 immunoreactivity from confocal images, we found that internalized hTDP43-GFP in microglia was more commonly observed in Clec7a KO mice than in WT mice (Fig. 4e). The orthogonal view of the single focal plane of hTDP43-GFP co-localization with Iba1 and CD68 immunoreactivity (magenta arrows, Fig. 4f, the magnified view of the box a in Fig. 4e) showed hTDP43-GFP fully surrounded by Iba1^+^CD68^+^ microglia. It also suggests that hTDP43-GFP was indeed inside CD68^+^ microglial lysosomes. In parallel, CD68 immunoreactivity in Clec7a-deficient microglia with internalized hTDP43-GFP (but not control GFP) was also significantly higher than in WT microglia (Extended Data Fig. 5i). Subsequent quantification showed a significantly increased percentage of microglia that contained hTDP43-GFP in Clec7a KO mice compared to WT mice (Fig. 4g) while control GFP alone was not found inside microglia, confirming that hTDP43, but not GFP, is the primary trigger to induce microglial phagocytosis. Consistently, a significantly higher percentage of hTDP43-GFP signals were observed inside Clec7a deficient microglia compared to WT microglia (Extended Data Fig. 5j). At the same time, microglia activation around hTDP43-GFP, indicated by Iba1 immunoreactivity, remained comparable in WT and Clec7a KO mice (Extended Data Fig. 5k). Additionally, we found that certain hTDP43-GFP signals co-localized with NeuN (white arrows, Fig. 4h i, v, vi) were phagocytosed by microglia (magenta arrows, Fig. 4h ii, iv), indicating that these neurons were stressed by hTDP43-GFP, may undergo degeneration, and were subsequently cleared by microglia. Overall, these results suggest that Clec7a-deficient microglia phagocytose and degrade hTDP43-GFP more actively than WT microglia.

Encouraged by these results, to directly visualize whether Clec7a deficiency alters microglial process dynamics to promote phagocytosis of hTDP43, we generated AAV9-hSyn-hTDP43-dTomato (dT) (Fig. 5a) and CX3CR1-eGFP^+^Clec7a^-/-^ mice to label microglia with the eGFP reporter for *in vivo* two-photon imaging of microglial processes (Fig. 5b). We first confirmed neuronal expression of hTDP43-dT by co-injecting a mixture of AAV-hSyn-hTDP43-dT and AAV-hSyn-GFP (to label neurons) in the motor cortex and observed predominant neuronal soma expression of hTDP43-dT (Fig. 5a) with hTDP43-dT signals also observed in GFP^+^ neurites (white arrows, Fig. 5a). Two-photon imaging was performed either at 5-minute intervals for 20 minutes or at 2-day intervals for 6 days, to specifically examine microglial process dynamics or phagocytosis of hTDP43-dT, respectively (Fig. 5b) following the cranial window surgery and AAV-hSyn-hTDP43-dT injection on the cortex of CX3CR1-eGFP^+^Clec7a^-/-^ and control CX3CR1-eGFP^+^ mice (∼P160). Active movement of eGFP^+^ microglial processes, either retracting (processes 1 and 2, P1-2, Fig. 5c i-iii) or extending (processes 3 and 4, P3-4, Fig. 5c iii-v), were commonly observed during time-lapse imaging (0’-20’, 5’ interval). By comparing process length at 0’ and 20’, microglial processes were further categorized as extension (20’-0’ length > 1μm), stable (-1μm < 20’-0’ length < 1μm), or retraction (20’-0’ length < -1μm), as shown in Extended Data Fig. 6a (based on 13 processes/4 microglia/1 mouse). Interestingly, a higher percentage of process extension was observed in Clec7a-deficient microglia in contact with hTDP43-dT while process retraction percentage remained essentially unchanged (Fig. 5d). Clec7a-deficient microglia in contact with hTDP43-dT also showed the highest process displacement frequency (Fig. 5e), calculated by averaging the total process displacement of individual microglia between consecutive imaging time points. On the other hand, Clec7a-deficient microglia without hTDP43-dT contact and WT microglia, regardless of hTDP43-dT contact, showed similarly lower process displacement frequency (Fig. 5e). Moreover, 20% of WT but no Clec7a-deficient microglia showed no process displacement (Extended Data Fig. 6b). These results suggest that Clec7a-deficient microglial processes become particularly active when in contact with hTDP43-dT, consistent with increased phagocytosis of hTDP43-GFP by Clec7a-deficient microglia observed above. In addition to microglial process dynamics in short imaging sessions, extended time-lapse imaging (from days 0 to 6) of eGFP^+^ microglia and hTDP43-dT also found examples that hTDP43-dT puncta (yellow arrows, Fig. 5f ii, iv) were actively cleared by eGFP^+^ microglia, confirming our immunohistology results above.

**Figure 5.**
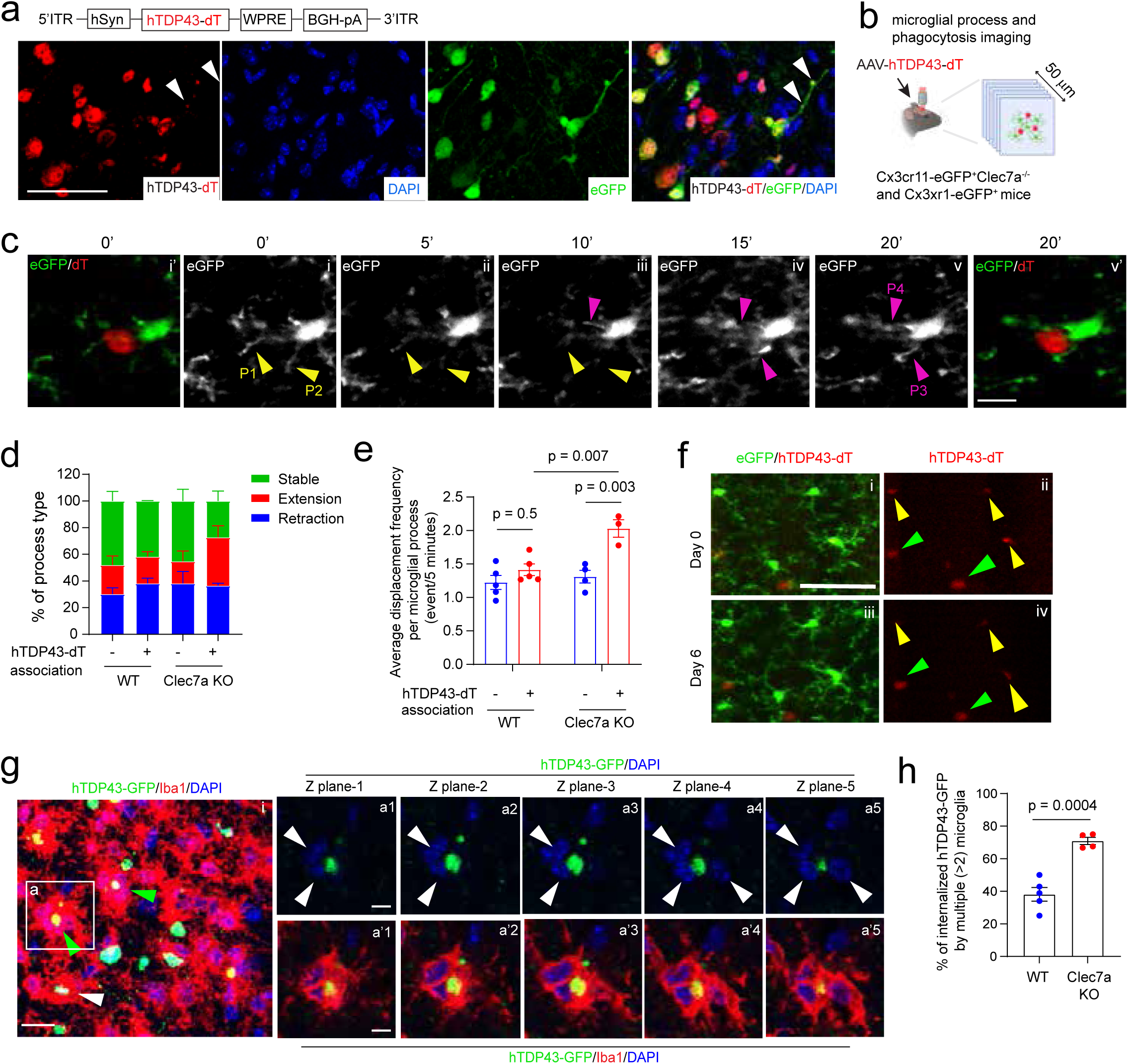
Clec7a deficiency promotes microglial process movement and microglial clustering in phagocytosing hTDP43. **a.** Design of neuron-specific hTDP43-dTomato (dT) AAV construct and representative images (63x) of co-expressed hTDP43-dT and eGFP in neurons followed by a focal injection of mixed AAV9-hSyn-eGFP and AAV9-hSyn-hTDP43-dT into the motor cortex of WT mice (P90). Scale bar: 50μm. **b.** Schematic view of *in vivo* two-photon imaging for monitoring microglial process dynamics and microglial phagocytosis of hTDP43-dT in AAV9-hSyn-hTDP43-dT injected Cx3xr1-eGFP^+^Clec7a^-/-^ and control Cx3xr1-eGFP^+^ unanesthetized mice. **c.** Representative time-lapse images of microglial process dynamics in contact with hTDP43-dT during a 20-minute live imaging period. i-v: monochrome images of eGFP^+^ microglial processes from 0 min (0’) to 20 min (20’). i’: multi-colored (eGFP^+^ microglia and hTDP43-dT) image same as the monochrome image at 0’; v’: multi-colored (eGFP^+^ microglia and hTDP43-dT) image same as the monochrome image at 20’. Yellow arrows (P1+P2): disappearing processes; purple arrows (P3+P4): emerging processes. Scale bar: 10μm. **d.** Percentage of microglial process types with or without contact with hTDP43-dT in the cortex of WT and Clec7a KO mice. **e.** Average displacement frequency per microglial process in WT and Clec7a-deficient microglia with or without contact with hTDP43-dT. n = 2-6 processes/microglia, 3-7 microglia/mouse, 3-5 mice/group. p values were determined by one-way ANOVA and post-hoc Tukey’s analysis. **f.** Representative *in vivo* imaging of eGFP^+^ microglial phagocytosis of hTDP43-dT at days 0 and 6, six weeks after AAV delivery and cranial window surgery in Cx3cr1-eGFP mice. yellow arrows: hTDP43-dT inside microglia; green arrows: hTDP43-dT^+^ signal with no apparent microglia nearby. Scale bar: 50μm. **g.** Representative images (40x) showing multiple microglia clustered around hTDP43-GFP in the motor cortex of Clec7a KO mice. a1-a5: hTDP43/DAPI from a series of Z-focal planes from the magnified view of the box a in i; a’1-a’5: hTDP43/Iba1/DAPI from the same series of Z-focal planes from the magnified view of the box a in i. Scale bar: 50μm (i), 20μm (a1-a5 and a’1-a’5). **h.** Quantification of the percentage of internalized hTDP43-GFP by multiple (≥ 2) microglia in the motor cortex of WT and Clec7a KO mice. n = 6 images from 3-4 sections/mouse, 4-5 mice/group. The p value was determined by unpaired Student’s t-test.

Additionally, it is evident that multiple (≥ 2) Clec7a-deficient microglia often simultaneously surround hTDP43-GFP for its clearance, indicated by multiple DAPI^+^ nuclei under same Iba1 immunoreactivity in consecutive individual Z focal planes (white arrows, a1-a5, a’1-a’5, magnified view of the box a in Fig. 5g i), while single microglial phagocytosis of hTDP43-GFP was also commonly observed (white arrows, Extended Data Fig. 6c i, iii). Subsequent quantification found that internalized hTDP43-GFP is much more commonly associated with multiple (≥ 2) microglia in Clec7a KO (∼70%) than in WT (40%) mice (Fig. 5h). This increased microglial clustering is not a result of increased microglial proliferation, as Ki67^+^Iba1^+^ microglia remained low and comparable in WT and Clec7a KO mice (Extended Data Fig. 6d). Similar multiple microglial clustering around hTDP43-dT was also observed (white arrows, Extended Data Fig. 6e). Together, our results suggest that Clec7a deficiency in microglia promotes microglial process dynamics and induces microglial clustering to facilitate microglial phagocytosis and clearance of hTDP43.

### Selective deletion of Clec7a in microglia attenuates motor neuron degeneration and delays disease progression in SOD1G93A ALS mice

Microglia have been previously shown to significantly impact disease progression in ALS^34^. How Clec7a-deficient microglia, with reduced neuroimmune and enhanced phagocytosis properties in ALS models as shown above, affect ALS disease progression is unknown. To begin unveiling the role of Clec7a in ALS pathogenesis, we monitored disease progression, defined by the duration from the peak weight to the end-stage (15s delayed righting reflex), of SOD1G93A^+^Clec7a^-/-^ and control SOD1G93A^+^ mice. The growth curve of mice in experimental groups is shown in Extended Data Fig. 7a. Excitingly, both SOD1G93A^+^Clec7a^-/-^ female and male mice have extended disease progression compared to SOD1G93A mice alone (Fig. 6a). The overall disease duration (from peak weight to the end-stage), also showed 13- and 12-days delay in SOD1G93A^+^Clec7a^-/-^ male and female mice (Fig. 6b), respectively. Clec7a is also expressed in several peripheral immune cell types, which can be activated in ALS conditions^35^. Particularly, peripheral sciatic nerve (SN) macrophages were shown to impact spinal cord microglial activation in SOD1G93A mice^36^. We found that Clec7a expression levels were up-regulated in SN macrophages in SOD1G93A mice (Extended Data Fig. 7b-c). The number of activated (CD68^+^) SN macrophages was also slightly increased in SOD1G93A^+^Clec7a^-/-^ compared to SOD1G93A^+^ mice (Extended Data Fig. 7d-e). Whether Clec7a increase in SN macrophages affects microglia activation and disease progression in ALS is unknown. To exclude that possibility, we generated Clec7a^f/f^ mice in which the exons 2 and 3 are floxed and can be deleted (Fig. 6c) when Cre is present. The Clec7a^f/f^ mice can be readily identified by PCR to recognize the WT and floxed allele (Extended Data Fig. 8a). The deletion of Clec7a exons 2 and 3 results in a frame-shifted truncated Clec7a and its non-sense mediated decay (NMD). By breeding with Clec7a^f/f^, SOD1G93A, and Cx3cr1-CreER mice, we generated microglia selective Clec7a deletion in the SOD1G93A model (M-Clec7a cKO/SOD1, SOD1G93A^+^Cx3cr1-CreER^+^Clec7a^f/f^) and littermate control (SOD1G93A^+^Cx3cr1-CreER^+^Clec7a^+/+^) mice. Both male and female M-Clec7a cKO/SOD1 mice showed similar growth trajectory as control SOD1G93A mice (Extended Data Fig. 8b). The overall experimental procedures and timeline are summarized in Fig. 6d. Tamoxifen (TAM, i.p., 75 mg/kg for 5 days) administration at P70 gives enough time (3 weeks) before disease onset (∼P90) for recombinant peripheral monocytes to turn over and to be replenished with non-recombined bone marrow progenitors, thus not affecting clec7a expression in the peripheral immune cells. We first confirmed that Clec7a expression is vastly reduced in microglia, indicated by 90% reduction of Clec7a^+^Iba1^+^ microglia numbers in spinal cords of diseased (P150) M-Clec7a cKO/SOD1 mice (Fig. 6e-f), without affecting Clec7a^+^Iba1^+^ macrophage number and Clec7a expression levels in SN macrophages (Extended Data Fig. 8c-e) of same M-Clec7a cKO/SOD1 mice following TAM administration. To examine whether selective deletion of Clec7a affects microgliosis and neuroinflammation in SOD1G93A mice, Iba1^+^ microglia numbers were quantified and showed significant reduction in spinal cords of M-Clec7a cKO/SOD1 relative to littermate SOD1G93A mice at the mid-disease (Fig. 6g-h). Pro-inflammatory cytokine IL-1β (Fig. 6i) was also significantly reduced in M-Clec7a cKO/SOD1 relative to littermate SOD1G93A mice at the disease end-stage, suggesting that IL-1β increase is particularly a result of microglial Clec7a activation in disease. To examine MN degeneration in M-Clec7a cKO/SOD1 mice, remaining MN numbers were quantified at the mid-disease stage based on choline acetyltransferase (ChAT) immunostaining and neuronal soma size (> 25 μm) (Fig. 6j) on spinal cord sections of control, SOD1G93A^+^, and M-Clec7a cKO/SOD1 littermate mice. While 50% of MNs were degenerated in SOD1G93A compared to WT mice (Fig. 6k), only 25% of MNs, on average, were degenerated in M-Clec7a cKO/SOD1 compared to WT mice at the mid-disease stage (Fig. 6k). Consistently, both male and female M-Clec7a cKO/SOD1 mice exhibited significantly slower progression compared to littermate SOD1G93A control mice, as shown in the Kaplan–Meier survival curve (Fig. 6l). The overall disease duration, defined from peak weight to end-stage, also showed 15- and 13-days delay in male and female M-Clec7a cKO/SOD1 mice, respectively (Fig. 6m).

**Figure 6.**
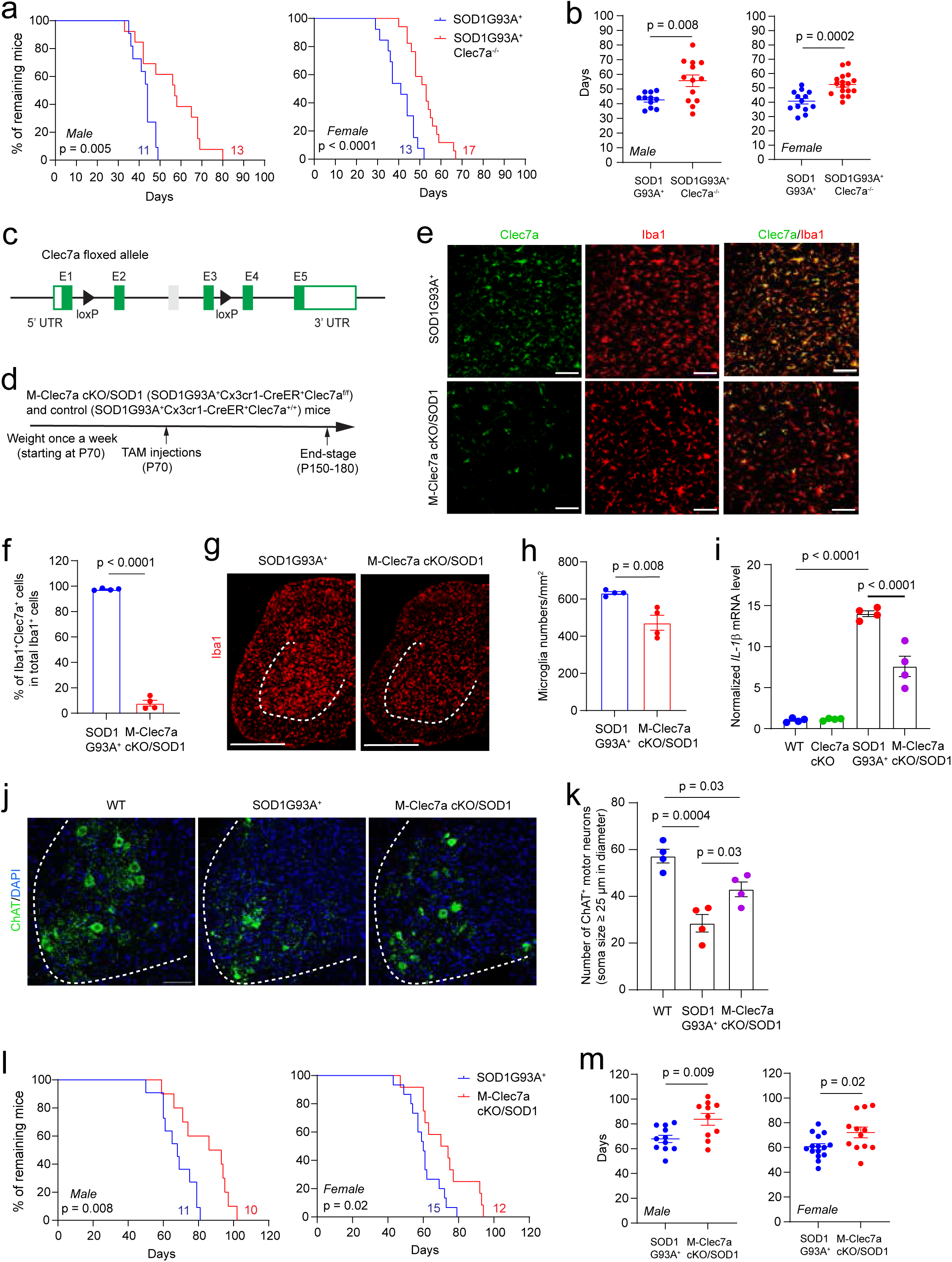
Selective deletion of Clec7a in microglia attenuates motor neuron degeneration and delays disease progression in SOD1G93A ALS mice. Kaplan-Meier plot (**a**) of disease progression and overall disease duration days (**b**) from peak weight to the end-stage in male and female SOD1G93A^+^Clec7a^-/-^ and control SOD1G93A mice. n = 11-13mice/group (male); n = 13-17 mice/group (female). p values determined by the Log-rank test in **a** and unpaired Student’s t-test in **b**. **c.** Schematic diagram of generating Clec7a floxed allele in Clec7a^f/f^ mice. E1-E5, exons 1-5; **d.** Experimental timeline of tamoxifen (TAM, 75 mg/kg, 5 days) injections and monitoring of disease progression until end-stage on M-Clec7a cKO/SOD1 (SOD1G93A^+^Cx3cr1-CreER^+^Clec7a^f/f^) and control (SOD1G93A^+^Cx3cr1-CreER^+^Clec7a^+/+^) mice. Note that SOD1G93A mice in C57BL/6J background (JAX stock #: 004435) were used in breeding to match the genetic background of Clec7a^f/f^ and Cx3cr1-CreER^+^ mice. Only heterogeneous Cx3cr1-Cre^+^ mice were used in experiments to avoid Cx3cr1 deficiency due to the Cre replacement. Representative Clec7 and Iba1 immunostaining images (20x) (**e**) and quantification (**f**) of the percentage of Clec7a^+^Iba1^+^ over total Iba1^+^ microglia in spinal cords of M-Clec7a cKO/SOD1 and control SOD1G93A^+^ mice at disease end-stage. Scale bar: 100μm. n = 6 images from 3 sections/mouse, 4 mice/group. p value determined by unpaired Student’s t-test. Representative images (20x) (**g**) of Iba1 immunostaining and quantification (**h**) of Iba1 immunoreactivity in spinal cords of M-Clec7a cKO/SOD1 and control SOD1G93A^+^ mice. Scale bar: 500μm. n = 6 images from 3-6 sections/mouse, 4 mice/group. p values were determined by unpaired Student’s t-test. **i.** mRNA expression levels of pro-inflammatory cytokines *IL-1β* in WT, Clec7a KO, M-Clec7a cKO/SOD1 and control SOD1G93A^+^ mice at disease end-stage. N = 4 mice/group. p values were determined by one-way ANOVA and post-hoc Tukey’s analysis. Representative images (20x, **j**) of ChAT immunostaining and quantification (**k**) of large (> 25μm) ChAT^+^ motor neurons in spinal cord sections of WT, M-Clec7a cKO/SOD1 and control SOD1G93A^+^ mice at mid-disease stage (P140±5). Scale bar: 100μm. n = 10 images from 5 sections/mouse, 4 mice/group. p values determined by one-way ANOVA and post-hoc Tukey’s analysis. Kaplan-Meier plot (**l**) of disease progression and overall disease duration days (**m**) from peak weight to the end-stage in male and female M-Clec7a cKO/SOD1 and control SOD1G93A^+^ mice. n = 10-11 mice/group (male), 12-15 mice/group (female). p values determined by the Log-rank test in **l** and unpaired Student’s t-test in **m**.

## Discussion

In our current study, by analyzing molecular profile and phagocytosis behaviors of Clec7a-deficient microglia in ALS models, we showed that Clec7a deficiency reduced diseased-induced expression of a cluster of microglial neuroimmune genes and pro-inflammatory cytokines. On the other hand, Clec7a deficiency promoted microglial process dynamics and enhanced phagocytosis of TDP43 aggregates. Selective deletion of Clec7a further attenuated motor neuron degeneration in spinal cords and extended disease duration of SOD1G93A ALS mice. As Clec7a is consistently and selectively up-regulated in microglia in various ALS mouse models and human ALS^1,5^, these results suggest that pathological up-regulation of Clec7a in microglia actively regulates disease microglial functions, leading to exacerbation of disease progression in ALS.

Disease induced up-regulation of Clec7a in microglia in SOD1G93A ALS mice has been shown to be dependent upon the TREM2 to APOE pathway^7^, which also induces expression of a number of DAM genes that are involved in lysosomal, neuroimmune, and phagocytic pathways. Whether Clec7a influences DAM signature or neuroimmune gene expression was not previously explored. Our results from the current study show that Clec7a deletion down-regulates microglial neuroimmune genes (Clec7a-M-DEG) induced in SOD1G93A ALS mice, such as *Cd86*, *Lilrb4*, *Parp14*, and *Igsf6*, etc. that have been previously reported to mediate microglial neuroinflammation^37–39^ and their expression were inversely correlated to disease progression in human ALS^5^. Another study using Clec7a shRNA-mediated knockdown in Parkinson’s disease 6-OHDA rat model also found similar effect in reducing neuroinflammatory responses^40^. Some of these neuroimmune genes also overlap with DAM molecular signatures in other studies^7,10^. Thus, our results suggest that Clec7a may act as an important disease-induced receptor to regulate neuroimmune signaling, by inducing expression of neuroimmune genes, in microglia to influence ALS disease progression. On the other hand, whether Clec7a acts downstream of the TREM2-APOE pathway to further induce DAM gene expression remains to be investigated in the future. Our results showed no changes of *Trem2* and *Apoe* in SOD1G93A^+^Clec7a^-/-^ mice relative to SOD1G93A mice, reinforcing that Clec7a acts downstream but not upstream of the TREM2-APOE pathway. Surprisingly, although Clec7a activation classically induces SYK phosphorylation to trigger downstream NF-κB signaling to drive gene expression^11^, we found that only a few of Clec7a-M-DEG promoters are predicted to bind to NF-kB. Instead, half of Clec7a-M-DEG promoters is predicted to bind to microglial enriched transcription factor Mef2a. In parallel, we found no clear evidence that p-SYK is involved in downstream signaling of Clec7a in microglia of diseased SOD1G93A mice. It is possible that SYK phosphorylation is difficult to detect, future studies with genetic approaches to manipulate SYK may help clarify its role in ALS models. Previous studies showed that Mef2a is able to regulate inflammatory gene expression in macrophages^41^ but specific involvement in ALS and other neurodegeneration remains little known. We found that its expression was specifically induced in disease microglia subclusters in SOD1G93A mice but was reduced in disease microglia subclusters of SOD1G93A^+^Clec7a^-/-^ mice. Our results point to an alternative Clec7a-Mef2a axis to regulate microglial neuroimmune genes in SOD1G93A ALS mice. How Clec7a signals are transmitted to Mef2a in regulating microglial neuroimmune gene expression remains to be further investigated.

Formation of abnormal protein aggregates, encoded by disease-causing mutant genes *SOD1*, *TARDBP*, *FUS*, and *C9ORF72*, is a common pathological feature in ALS, which can significantly influence ALS disease progression^27,28^. In particular, both wild type and mutant TDP43 can be commonly depleted from nucleus and become hyperphosphorylated to form cytoplasmic inclusions in ALS and frontotemporal dementia (FTD)^31,33^. Microglia play a central role in phagocytosing and clearing abnormal TDP43 aggregates, either in degenerating neurons or secreted extracellularly^30,42^. Microglial clearance of neuronal TDP43 has been shown to be neuroprotective in TDP-43 ΔNLS (rNLS8) ALS mice in which hTDP-43 pathology could be reversibly induced in neurons^43^. Disease-induced microglial TREM2 has also been shown to directly interact with TDP43 to promote TDP43 phagocytosis^30^, which contributes to TREM2-mediated neuroprotection. Our current study shows Clec7a deficiency promotes microglia phagocytosis and degradation of (neuronal and extracellular) TDP43 aggregates. Consistently, our *in vivo* two-photon imaging also revealed enhanced microglia process dynamics in Clec7a-deficient microglia in contact with TDP43. Clec7a-deficient microglia were also more actively recruited to surround TDP43 aggregates for their phagocytosis and degradation. Our results are in contrast to recent reports that Clec7a promotes microglial phagocytosis of β-amyloid^44^. Whether β-amyloid and TDP43 aggregates directly interact with Clec7a for their phagocytosis remains unknown, but they do not possess the typical β-glucan structure recognized by the extracellular C-type lectin-like domain on Clec7a^11^. It is possible that the opposite effect of Clec7a on phagocytosis of β- amyloid (with cross-β sheet structure) and amorphous TDP43 aggregates may result from their structural differences. As a result, Clec7a signaling activation may recruit different microglial phagocytic receptors or induce selective cytoskeleton changes to mediate the phagocytosis of β-amyloid or TDP43 aggregates. Indeed, previous studies have shown that Clec7a collaborates with Toll-like receptor 2 (TLR2) to enhance recognition and clearance of microbes in dendritic cells and macrophages^45^. Clec7a can also act synergistically with complement receptor 3 (CR3) in macrophages in response to fungal infection^46^. Specific modulation of phagocytic receptor activity by Clec7a in microglia remains to be better understood in the future.

Microglial responses and modulatory roles in ALS disease progression are heterogeneous and complex. Selective depletion of proliferating microglia using CD11b targeted expression of mutant herpes simplex virus thymidine kinase (TK) had no effect on motor neuron degeneration in SOD1G93A mice^47^, but the efficiency of this strategy only resulted in 30-50% reduction of CD11b^+^ microglia^47^. Presymptomatic administration of CSFR1 inhibitor GW2580 substantially reduced microglia proliferation and significantly delayed disease progression in SOD1G93A mice^48^. On the other hand, CSFR1 antagonist PLX3397-mediated microglial depletion prevented early recovery of motor functions in rNLS8 ALS mice^43^, suggesting an important neuroprotective role for microglia. However, these studies provided limited mechanistic insights into microglia-mediated disease pathways. In contrast, our current study defines pathogenic roles of microglial Clec7a pathway in ALS disease progression, providing new insights into how microglia modulate ALS. Importantly, selective and induced deletion of Clec7a in microglia showed similar extended disease duration as germline Clec7a deletion in SOD1G93A mice, suggesting that particularly microglial Clec7a but not peripheral Clec7a contributes to disease progression. There are several important questions that remain to be answered, including how Clec7a is activated by disease-relevant ligands or patterns and what the downstream signaling is to mediate neuroimmune gene expression and phagocytosis behaviors. Answers to these questions will further advance our understanding of the microglial pathways in ALS pathogenesis.

## Acknowledgments

We thank Dr. Albert Tai and Tufts University Core Facility (TUCF) -Genomics for providing help in preparation of single cell sequencing library and sequencing. Imaging was performed with the assistance of the Tufts Center for Neuroscience Research. We thank Dr. Long-Jun Wu (University of Texas Health Science Center at Houston) for providing AAV-hTDP43-GFP virus. We thank Dr. Zuoshang Xu (University of Massachusetts Chan Medical School) for providing spinal cord tissues of PFN1C71G ALS mice. We thank Caroline Reynolds for providing initial help with two photon image analysis. We thank Target ALS Post-Mortem Consortium (University of California San Diego and Georgetown University) for providing human control and ALS spinal cord tissues. This work was supported by NIH grants RF1AG057882, RF1AG05961, R01NS125490, and R01AG078728 (YY).

## Author contributions

X.C. and Y.Y. designed the experiments and wrote the manuscript. X.C. performed most of the experiments. H.Y. helped with single cell preparation and scRNA-seq data analysis, tissue collection, immunostaining and image analysis. S.S. and M.P. performed surgeries and imaging for in vivo two photon experiments and wrote manuscript. J.H. performed stereotactic injections, tissue collection, immunostaining and data analysis. H.W. and J.W. performed scRNA-seq data analysis and wrote manuscript. V.V., E.K., and A.K. performed image analysis. H.T. performed image analysis, body weight recording, and genotyping. E.K. performed image analysis. T.S. provided Macro Scripts of ImageJ for image analysis.

## Data availability statement

All data supporting this study is available upon request. scRNA-seq data has been deposited to Gene Expression Omnibus (GEO) repository (accession number: GSE328178, secure token: gvstaucsvzobrcl) at the NCBI.

## Materials and Methods

### Animals

The Clec7a flox mice were generated by homologous recombination. The two loxp cassettes were subcloned into the Clec7a ATG-containing-exon targeting vector, at the location of the upstream intron of Exon2 and the downstream intron of Exon4 respectively, to generate the final loxp flanked Exon2-4 targeting vector. The targeting vectors were linearized and transfected into the Embryonic Stem (ES) cell line to induce homolog recombination. G418-resistant ES clones were screened by Southern blot analysis and were subsequently injected into C57BL/6J blastocysts to obtain chimeric mice following standard procedures. Both ES cell transfections and blastocyst injections were performed by Biocytogen (Worcester, MA). Chimeric mice were bred with C57BL/6J mice to obtain germline transmission. The wild type (WT) C57BL/6J (Stock No: 000664), B6.SOD1-G93A (Stock No: 004435), B6SJL.SOD1-G93A (Stock No: 002726), Clec7a KO (Stock No: 012337), Cx3cr1CreER knock-in (Stock No: 021160) and CX3CR-1GFP knock-in (Stock No: 005582) mice were all obtained from The Jackson Laboratory. Clec7a KO mice are normal without obvious adverse phenotypes including motor deficits, as previously described^13^ and also confirmed by our own observations. Both male and female mice were used in all experiments. All mice were maintained on a 12 h light/dark cycle with food and water ad libitum. Care and treatment of animals in all procedures strictly followed the NIH Guide for the Care and Use of Laboratory Animals and the Guidelines for the Use of Animals in Neuroscience Research. Animal protocols used in this study have been approved by Tufts University IACUC committee.

### Drug administration

Tamoxifen (Sigma-Aldrich, CAS# 10540-29-1) was dissolved in Corn oil (Sigma-Aldrich, CAS# 8001-30-7) at a concentration of 20mg/ml. TAM (75mg/kg) was administrated to experimental mice (10 weeks) by intraperitoneal (i.p.) injections for 5 consecutive days.

### Human samples and DAB Immunohistochemistry

Postmortem spinal cord paraffin sections were applied from Target ALS human postmortem tissue cores at University of California San Diego (UCSD) and Georgetown University. Total 8 cases of sALS (age 59 ± 10.6 years (mean ± s.d.)), 6 cases of fALS (age 67± 8.4 years) and 9 cases of non-neurological disorder control (age 70± 13.8 years). The study was approved by the Tufts Biospecimen Committee. For DAB staining, paraffin embedded human sections were first incubated at 65°C for 30min to facilitate wax removing steps. The slides undergo deparaffinization through washing in graded xylene and ethanol series (100% Xylene twice, 50% Xylene and 50%Ethanol mixture, 100%, 90%, 70%, and 50% Ethanol). Antigen retrieval was performed by incubating slides in Citrate buffer PH6.0 at 100°C for 30min. Human sections were rinsed three times in PBS and incubated in 3% hydrogen peroxide for 20min to quench endogenous peroxidase activity. Immunohistochemistry staining for primary antibody anti-human Clec7a (MAB1859, mouse, R&D systems, 1:50) were performed using a Vecta stain-ABC universal plus kit (Cat.NO. PK-8200, Vector Laboratories) following the manufacturer’s instructions. After DAB staining, sections were further counterstained with Harris Hemotoxylin (StatLab) for 2min and washed under tap water. Subsequently, counterstained sections were dehydrated through 50%, 70%, 90% and 100% Ethanol solutions. After immersing in 100% Xylene twice. Coverslips were placed on slides using DPX (Sigma-Aldrich) mounting medium for observation. Brightfield images were captured with light microscope (Olympus CX41), using Stream Essentials software to process.

### Stereotaxic intracerebral injection of AAVs

WT and Clec7a KO mice (8-10 weeks of age) were anesthetized with 2% isoflurane. AAV9-CAG-hTDP43-GFP (0.5μl, 0.9 x10^13^ gc/ml) or AAV9-CAG-GFP (0.5μl, 1.0 x10^13^ gc/ml) were stereotaxically injected into the right or the left primary motor cortex respectively (Bregma -1.4, *X* ±1.5mm, *Z* 0.5mm). AAV9-CAG-hTDP43-GFP virus was obtained from Long-Jun Wu lab (University of Texas Health Science Center at Houston). The microinjections were performed at a rate of 0.05μl/min using an automated stereotactic injection apparatus (UMP3 with MICRO2T, WPI). For neuronal hTDP43-dT and eGFP expression, AAV9-hSyn-hTDP43-dTomato/AAV9-hSyn-eGFP virus mixture (0.6μl, 3.5 x10^10^ gc/3.5 x10^9^ gc) was stereotaxically injected into the right primary motor cortex of experimental mice (Bregma - 1.4, *X* ±1.5mm, *Z* 0.5mm). pAAV-hSyn-hTDP43-dTomato (dT) plasmid was produced by Vector Builder and AAV virus was packaged at Boston Children’s Hospital Viral Core. AAV9-CAG-GFP (#37825) and AAV8-hSyn-EGFP (#50456) virus were purchased from Addgene. Post-operative care included injections of buprenorphine according to the IACUC requirement. Animals were perfused 4 weeks after injections.

### *In vivo* two photon surgeries, imaging, and analysis

All *in vivo* imaging procedures were conducted in accordance with guidelines and protocols of the University of Texas Health Science Center at San Antonio (UTHSCSA) Institutional Animal Care and Use Committee (protocol number 20140023AR). Mice were kept in the Laboratory Animal Resources facility with ambient 72–78 °F temperature and 30–70% humidity and had ad libitum access to water and chow. Mice were maintained on a reverse 12h-light/12h-dark schedule (lights off at 10 a.m., on at 10 p.m.), and most experimental procedures were completed during the dark cycle.

For stereotaxic delivery of AAV9-hSyn-hTDP43-dTomato in the left somatosensory cortex, mice at 16-18 weeks were anesthetized with 100 mg/kg ketamine (Hikma Pharmaceuticals USA Inc.) and 10 mg/kg xylazine (AnaSed from VETone) administered i.p. and positioned on a heating pad to maintain body temperature at ∼36 °C. Hair was removed and skin disinfected using povidone-iodine. Before skin incision, 5 mg/kg meloxicam (Covetrus) was administered s.c. Skin and muscles were removed from the skull, and 3% hydrogen peroxide was applied to disinfect and prevent bleeding. The periosteum was shaved off, and remaining muscle surrounding the exposed skull was covered with a thin layer of cyanoacrylate cement. Following installation of a burr hole at coordinates anteroposterior (AP) −1.2 to −1.7 mm and mediolateral (ML) 2.7 to 3.3 mm relative to bregma, AAV was injected at a depth of 1.5 mm from the cortical surface, with a total volume of 0.5 µL (6.9 x10^13^ gc/ml) within 10 minutes. Immediately following viral injection, a stainless-steel head plate with a 4 mm in diameter central opening was mounted over the left somatosensory cortex and secured using dental cement (C&B Metabond, Parkell Inc., Brentwood). Mice were placed in a heated cage for recovery. 15 days later, animals underwent a second surgical procedure, during which a cranial window was implanted in the center of the head plate opening. Under ketamine/xylazine anesthesia and on a heating pad, mice underwent a craniotomy where a 2 mm x 2 mm portion of skull was removed. Dura mater was torn and pushed aside, and the exposed brain tissue was covered with a glass window comprised of three fused layers of No. 1 Corning cover glass. The window edges were sealed to the skull with dental cement (Ortho-Jet-Acrylic-Powder, Lang) with a small amount of pooled CSF preventing direct contact of dental cement with brain tissue. After 1 week of post-surgery recovery, mice were habituated to head restraining.

All *in vivo* two photon imaging was performed using an unanesthetized, head-restrained mouse on a treadmill, while mouse activity was not monitored. A resonant scanning version of the MOM (Sutter Instruments) was used with a x16, 0.80 NA water-immersion objective (Nikon). A pulsed Ti:Sapphire laser beam at 80 MHz repetition rate, <140 fs pulse width (Coherent Inc., Chameleon Ultra II) was tuned to 1000 nm to achieve simultaneous two-photon excitation of GFP and dTomato. To minimize brain injury, laser power was attenuated to 20-30 mW at the front aperture of the objective. Emitted light was detected using two photomultiplier tubes (H10770PA-40; Hamamatsu Photonics) respectively equipped with filters to separate the GFP from the dTomato signals. The awake mouse was placed on a custom-made linear treadmill, and the head plate was secured under the microscope objective. Data were acquired using volumetric scanning with a step size of 1 µm, 50 slices per volume covering approximately 50 µm in the z-plane, and at a rate of 30 slices per second and 0.59 volumes per second. Acquired slices were 200 µm x 200 µm at a resolution of 1024 pixels per line and 512 lines per slice. The microscope was controlled by an Xi Computer Corporation personal computer (Intel(R) Core (TM) i7-5930K CPU @ 3.50 GHz, 16 GB of RAM) running ScanImage (v5.0; Vidrio Technologies, LLC) software within MATLAB R2019a (MathWorks). Ten volume scans were acquired consecutively as a block, volume scans appearing blurred due to animal motion were removed and remaining unperturbed volume scans were averaged for improved resolution of fine structure. For the quantification of microglia motility, imaging blocks acquired at 5 minutes intervals were compared. For the quantification of dynamic interactions of microglia with their environment including hTDP43-dTomato molecules, imaging blocks of respective fields of view (FOVs) were acquired every other day. Up to four FOVs were analyzed per mouse.

For microglial process dynamics analysis, all *in vivo* two photon images of microglial motility assay were processed using Image J software (National Institutes of Health, Bethesda, MD). 50 sections from 1 z-stack (50 slices from 1 volume) were average projected to generate a single image of each volume. To neutralize the artifact movements in obtained images, 10 sequential z-stacks of images (volumes) were first subjected to the ImageJ plug-in StackReg/TurboReg with Rigid Body mode, and the aligned z-stacks were maximally projected to a single image at every frame. For each FOV, five frames acquired by time-lapse imaging within 5 min interval (Time 0’, 5’, 10’, 15’ and 20’) were further aligned with plug-in StackReg/TurboReg using Time 0’ image as reference. Then, a maximum intensity projection image (MPI) was computed from the stack of five registered frames. On this MPI image, microglial processes-paths were manually traced, from soma to the tip of the process. The image intensity along each path was plotted for each imaged time points. Length of a process-path was measured by the sum of selected bin numbers that had intensity values above 2, a bin indicating 0.195 μm in length according to the scale. Moving frequency of each process was calculated by number of displacements occurred during recording period, with each displacement having more than 1μm length change relative to previous time point. The individual microglial movement was determined by the observation of the displacement of at least one process between end and start time points.

### Primary cortical microglia culture and immunocytochemistry staining

Primary cortical microglial cultures were prepared from P1-3 mouse pups. Cerebral cortices were dissected, and meninges were removed in cold HBSS buffer. Tissues were digested with 0.05% trypsin-EDTA solution for 10mins in a 37°C water bath. The enzymatic reaction was ceased by adding 10%FBS containing DMEM. Tissues were then triturated with fire polished Pasteur pipettes to obtain single cell preparation. Cell suspension was filtered through a 70μm strainer and cell pellets were collected by centrifugation at 1,000g for 5min. Cell density was determined using hemocytometer, and cells were seeded in Poly-L-lysine (PLL) coated 10cm culture dishes at a density of 3.0 x10^6^ cells/dish with glia culture medium (DMEM supplemented with 10%FBS and 1%Pen-Strep). The culture medium was refreshed the next day to remove cell debris and then change culture medium every 5 days. In 9-12 days, the glial culture formed a confluent cell layer with microglia growing on top. To isolate microglia, dishes were vigorously tapped on the bench top. Floating microglia were collected from conditioned culture medium, then were seeded on PLL coating coverslips in 24-well plate (5.0 x10^4^ cells/well) with culture medium containing half conditioned medium. Microglia were ready for experiment the following day. Primary WT microglia cultured in coverslips were treated with LPS 10μg/ml (Lipopolysaccharides from Escherichia coli O55:B5, Sigma-Aldrich) for 2hr at 37 °C. Cells were washed with PBS and fixed using 4%PFA for 15min before immunocytochemistry staining. Cells were permeabilized with 0.2% Triton X-100 for 10 min and blocked in 5% bovine serum albumin for 1hr and incubated with the following primary antibodies overnight at 4°C: anti-phosopho-SYK (Y352) (#2717, Cell Signaling, rabbit, 1:200), anti-Iba1 (234011, mouse, Synaptic Systems, 1:100). After incubation with the primary antibodies, cells were washed three times with PBS, incubated with following secondary antibodies for 1 h at room temperature: Donkey Alexa Fluor 488 anti-rabbit and Goat Alexa Fluor 633 anti-mouse (1:1000, Invitrogen), and mounted with Prolong^TM^ Gold antifade reagent with DAPI (Invitrogen). Cells were imaged using Lecia FALCON confocal laser scanning microscope (7-12μm, Z stack with 1.0μm step), magnified with 63X oil objectives. Images are processed with LAS X software.

### Immunostaining, imaging, and analysis

Tissues were prepared as previously reported (Yuqing, et al. 2019). Mice were deeply anesthetized by Ketamine (100mg/kg) + Xylazine (10mg/kg) in saline by intraperitoneal (IP) injection and perfused intracardially with 4%paraformaldehyde (PFA) in PBS. Tissues including spinal cord, brain, sciatic nerve, were dissected and kept in 4%PFA overnight at 4°C, then cryoprotected by immersion in 30% sucrose for 48h. Tissues were embedded and frozen in OCT-Compund Tissue-Tek® (Sakura, Tokyo Japan). Coronal sections (20μm) of spinal cord and brain, Longitudinal sections of sciatic nerve (12 μm) were prepared with a cryostat (Lecia HM525) and mounted on glass SuperFrost ^+^ slides (Fisher Scientific). Slides were rinsed three times in 0.2% Triton-X-100 PBS (0.2% PBST), followed by blocking with 10% donkey serum and 0.2% PBST for 1h at room temperature before applying the primary antibody diluted in 10% block buffer overnight at room temperature. Primary antibodies used here include anti-Clec7a (R1-8G7, rat, InvivoGen, 1:200), anti-Iba1 (019-19741, rabbit, FujiFilm, 1:500), anti-Iba1 (NB100-1028, goat, Novus, 1:50), anti-Cd206 (AF2535, goat, R&D systems, 1:100), anti-Mef2a (sc-17785, mouse, Santa Cruz, 1:50), anti-ChAT (AP144-P, goat, Millipore, 1:50), anti-NeuN (MAB377, mouse, Millipore, 1:400), anti-GFAP (16825-1-AP, rabbit, Proteintech, 1:2000), anti-GFAP (75-240, mouse, UC Davis/NeuroMab,1:100), anti-Olig2 (AB9610, rabbit, Millipore, 1:200), anti-Cd68 (ab237968, rat, Abcam, 1:200), anti-Phosopho-TDP43 (Ser409/410) (26H10, rabbit, DSHB, 1:500) and anti-Ki67 (NB600-1252, rabbit, Novus, 1:100). For the staining of anti-human TDP43 (60019-2-Ig, mouse, Proteintech, 1:100), antigen retrieval was performed ahead of staining, using Sodium Citrate buffer (10 mM Sodium citrate, 0.05% Tween 20, pH 6.0) heating at 65°C for 5mins. For the staining of anti-P62/SQSTM1 (H00008878-M01, mouse, Novus, 1:50), primary antibodies were incubated for 48hr at 4°C. After washing slides three times in 0.2% PBST, corresponding secondary antibodies Donkey Alexa Fluor 488 anti-rabbit, -mouse, -goat, Donkey Alexa Fluor 555 anti-rabbit, -mouse, Donkey Alexa Fluor 647 anti-goat, Goat Alexa Fluor anti-rat (Invitrogen, 1:1000), were added for 1h at room temperature. The sections were rinsed three times in 0.2% PBST before mounting. All the stained slides were mounted with Prolong^TM^ Gold antifade reagent with or without DAPI (Invitrogen).

Sections of brain and spinal cord were imaged using Lecia FALCON confocal laser scanning microscope (15-20 μm, Z stack with 1.0 μm step), images were processed with LAS X software. Images of SN sections were taken by Zeiss Axio fluorescence microscope, using ZEN2 software to process. Six to ten diverse regions across each sample were imaged. Cells of interest were counted, fluorescence signal intensity and colocalization were quantified using ImageJ software.

For the quantification of Mef2a expression in Iba1^+^ microglia at Mid-disease stage of SOD1G93A and SOD1G93A/Clec7a KO mice, the ROI of Iba1^+^ Dapi^+^ microglial soma was manually selected using ImageJ “Oval” tool, and the “Integrated Density (IntDen)” of Mef2a staining in soma was measured using selected ROI from maximum projection image, then the average IntDen of Mef2a per microglial soma in each sample was analyzed.

For the quantification of microglial phagocytosis of hTDP43-GFP, the proportion of hTDP43-GFP^+^ Iba1^+^ microglia among Iba1^+^ microglia, and the ratio of hTDP43-GFP^+^ NeuN^+^ Iba1^+^ microglia in hTDP43-GFP^+^ Iba1^+^ microglia were calculated by the numbers of corresponding cell populations counted using Image J “Cell Counter”, based on single plane images from stacks; The number of hTDP43-GFP puncta in imaged field was measured by ”Analyze Particles” (Size > 5 μm^2^) based on maximum projection image, and ROIs of hTDP43-GFP puncta were used to select hTDP43-GFP puncta colocalizing with Iba1^+^ microglia in single plane images, the following ratio of hTDP43-GFP internalized by microglia was calculated accordingly.

For the quantification of motor neurons in the lumbar region (L3-L5) of spinal cord, total 60 consecutive sections were collected. Counting of motor neurons was performed at the ventral horn in every tenth section for five sections in total per animal. The images of ChAT staining were acquired at x 20 magnification. ChAT^+^ motor neurons were analyzed using Image J “Cell Counter” with clearly visible nucleus and soma size ≥ 25 μm in diameter, to avoid small ChAT^+^ interneurons or fragmented cell debris. The observer was blinded to the genotype of studied mice.

### Immunoblotting

Primary microglial pellets were homogenized with lysis buffer (Tris-HCL pH 7.4, 20 mM, NaCl 140 mM, EDTA 1 mM, SDS 0.1%, Triton-X 1%, Glycerol 10%). Protein inhibitor cocktail (P8340, Sigma) and phosphatase inhibitor cocktail 3 (P0044, Sigma) was added in a 1:100 dilution to lysis buffer prior to tissue homogenization. Total protein amount was determined by DC™ Protein Assay Kit II (Bio-Rad), then lysates were loaded on 4-15% Mini-PROTEAN TGX Stain-Free Protein Gels (Bio-Rad). Separated proteins were transferred onto a PVDF membrane (Bio-Rad) with Trans-Blot Turbo Transfer System (Bio-Rad). The membrane was blocked with 5% fat-free milk in TBST (Tris buffer saline with 0.05% Tween 20), then incubated with primary antibody anti-Clec7a (23042, rabbit, Cell signaling, 1:500), anti-human-SOD1 (2770, rabbit, Cell signaling, 1:500), and anti-β-actin (A1978, mouse, Sigma, 1:2000) overnight at 4°C. The next day, Secondary antibodies, anti-mouse IgG-HRP (NA931, GE Healthcare, 1:10000), anti-rabbit IgG-HRP (NA934, GE Healthcare, 1:5000), were incubated for 1hr. Bands were visualized on Chemidoc MP imaging system (Bio-Rad) with Clarity Western ECL substrate (Bio-Rad). Immunoblots were analyzed using ImageJ.

### Preparation of single-cell suspension and single-cell RNA sequencing

The whole spinal cord (SC) tissue was harvest and dissociated. After anesthesia, mice were transcardially perfused with cold PBS, SCs were extracted from the spines by flushing with PBS. SCs dissociation were performed using a Neural tissue dissociation kit (Miltenyi Biotec, #130-092-628) following the manufacturer’s instructions with some modifications: 1) Add Enzyme mix 1 (100μl Enzyme P in 3800 μl Buffer X) and Enzyme mix 2 (20μl Enzyme A in 40 μl Buffer Y) to each spinal cord in order, incubate it in 37°C water bath with gently inverting every 3-5 min to resuspend settled tissues. 2) Dissociate tissue mechanically using the wide-tipped, fire-polished Pasteur pipette in decreasing diameter. 3) Filter the single-cell suspension through a 70μm cell strainer. 4) Cells were resuspended in cold PBS supplemented with 0.5% BSA containing Myelin Removal Beads II (Miltenyi Biotec, #130-096-733) to remove myelin, following the manufacturer’s protocol. After myelin removal, Cell number and viability were measured by TC20 Automated Cell Counter (Bio-Rad). The single-cell suspensions were kept on ice for no longer than 30min until further processing.

For the scRNA-seq experiments, total 14 spinal cord samples from WT (n=3), Clec7aKO (n=4), SOD1G93A (n=3) and SOD1G93A/Clec7a KO (n=4) mice were processed in four separate days. For each sample, cells were first diluted at a density of 1,000 cells/μl, then 10,000 cells were loaded into a Chromium NextGEM Chip G (10x Genomics) and processed following the manufacturer’s instructions. Then, scRNA-seq libraries were prepared with the Chromium Single Cell 3’ Kit v3.1 (10x Genomics) following the manufacturer’s instructions. For each sequencing run, eight FASTQ files were generated, corresponding to Index 1 (I1), Index 2 (I2), Read 1 (R1), and Read 2 (R2) across two lanes (L001 and L002). The run employed a paired-end, dual-index configuration on NextSeq 550 instrument (Illumina) platform, producing four reads per cluster: I1 (10 bp) and I2 (10 bp) for sample demultiplexing, R1 (28 bp) containing the cell barcode and unique molecular identifier (UMI), and R2 (90 bp) capturing the cDNA insert for transcript alignment and gene expression quantification.

### Single-cell RNA sequencing data analysis

Raw single-cell RNA sequencing reads were first processed individually for each sample using the 10x Genomics CellRanger software^49^ suited with default parameters, and aligned to the mouse reference genome (mm10). Following alignment and gene counting, data from all samples were imported into the Seurat R package^50^ for downstream analysis and subsequently merged into a single dataset. To ensure data quality, stringent filtering criteria were applied. Cells with fewer than 300 detected genes were removed to exclude low-quality or empty droplets, while cells with more than 7,000 genes were discarded to minimize the inclusion of doublets. In addition, genes expressed in fewer than three cells across the dataset were excluded. To control for artifacts associated with stressed or dying cells, we further removed cells in which mitochondrial transcripts represented more than 10% of total expression. Finally, mitochondrial genes themselves were excluded from the expression matrix to reduce confounding signals.

Normalization was performed in Seurat using default log-normalization parameters. To address batch variation across samples, we applied the FindIntegrationAnchors function followed by IntegrateData, thereby aligning datasets into a shared expression space. Dimensionality reduction was then carried out with principal component analysis (PCA), retaining the top 30 principal components for downstream clustering. Cell clustering was performed using the FindClusters function at a resolution of 0.1, and results were visualized with Uniform Manifold Approximation and Projection (UMAP). Clusters were annotated based on well-established cell-type marker genes, enabling robust identification of major cell populations. Specific cell types, including microglia, astrocyte, neuron, and endothelial cells, were further isolated using Seurat’s subset function. Within cell clusters, differentially expressed genes (DEGs) were identified using FindAllMarkers, applying default thresholds for fold change and adjusted significance. To investigate biological pathways associated with transcriptional changes, gene set enrichment analysis (GSEA) was performed using the fgsea R package (v1.16.0). Enrichment was tested across all MSigDB C5 gene ontology signature sets, with 1,000 random permutations used to assess significance. Gene sets achieving a nominal p value < 0.05 were considered significantly enriched.

For complementary pseudo bulk RNA-seq analysis, raw gene counts were extracted from the Seurat object (RNA assay) and aggregated by sample group and cell type, producing a gene-by-sample count matrix that reflects average expression across groups. The resulting matrix was processed with DESeq2, using size factor estimation and standard normalization procedures, thereby generating normalized expression profiles comparable to those obtained from conventional bulk RNA-seq. These pseudobulk values were used for downstream statistical testing and integrative comparisons. Biomolecular signal pathways were identified through QIAGEN Ingenuity Pathway Analysis (IPA) platform.

### RNA isolation and quantitative PCR

Mouse spinal cord tissues were freshly collected and immediately frozen on dry ice after perfusion with cold PBS. Total RNA of tissue was extracted with TRIzol reagent and treated with DNase I, cDNA was then synthesized using SuperScript III First-Strand Synthesis System according to the manufacturer’s instructions (Invitrogen/Thermo Fisher Scientific). Reverse transcription products were amplified with the StepOnePlus Real-Time PCR System (Life Technology) using SYBR Green PCR Master Mix (Applied Biosystems) following the manufacturer’s instructions. Total 100ng cDNA of each sample was used for qPCR reactions. Data was normalized to the level of *GAPDH* in each individual sample. The 2^-ϪϪCt^ method was used to calculate relative expression changes. The primers used in the qPCR were *IL-1β*, 5’-GCAACTGTTCCTGAACTCAACT-3’ (Forward) and 5’- ATCTTTTGGGGTCCGTCAACT-3’ (Reverse); *TNF-α*, 5’- GGAACACGTCGTGGGATAATG-3’ (Forward) and 5’- GGCAGACTTTGGATGCTTCTT-3’ (Reverse); *GAPDH*, 5’- AGGTCGGTGTGAACGGATTTG-3’ (Forward) and 5’- TGTAGACCATGTAGTTGAGGTCA-3’ (Reverse).

### Monitoring of body weight and determination of end-stage

Body weights were performed every five days for the Clec7a KO mice cohort or weekly for the Clec7a cKO mice cohort during the noon time, beginning at postnatal (P) day 60 or 70 until the death of the animals. The survival of mice was calculated from birth to the end point. Disease end point in mice was determined by the loss of righting reflex in 15s when placed on their side.

### Statistical analysis

All statistical analysis were performed, and graphs were generated using GraphPad Prism 10. Experimental sample sizes and statistic tests were described in the figure legends. Results of analysis were presented as Mean ± S.E.M. Group differences in each assay were either analyzed by two-tailed Student’s t-test in comparison of two groups or analyzed by one-way ANOVA in comparison of multiple groups. Survival curves were analyzed using a log-rank (Mantel-Cox) test. Statistical significance was tested at a 95% (p < 0.05) confidence level and p values are shown in each graph. Mice were grouped according to genotype before randomly assigned to the experimental groups. The investigators were blinded to group allocation during data collection and analysis. Animals or samples were excluded from analysis in stance of technique failure. Outliners in each experimental group in disease progression were removed based on Mean ± 1.5 Standard Deviation (SD). No data was excluded for other reasons. No custom code was used in the analysis.

