## supplementary figures for "Genetic suppression of myeloid receptor Clec7a attenuates microglia neuroinflammation and promotes microglial phagocytosis to delay disease progression in ALS models"

Chen X et al., 2026

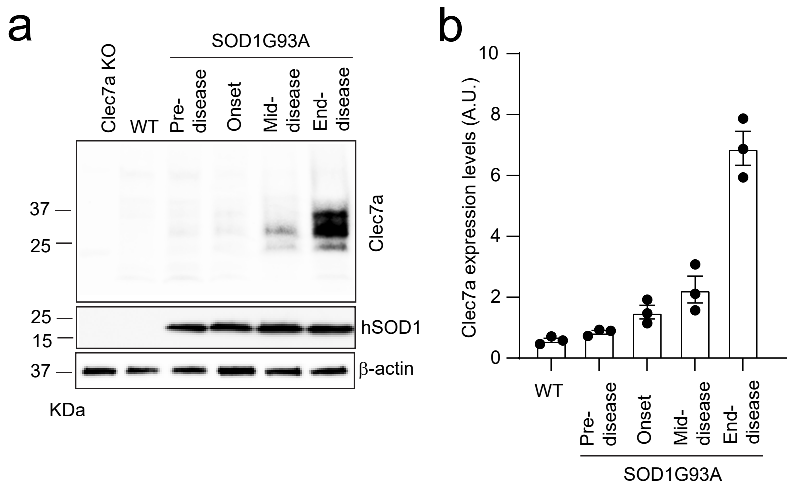


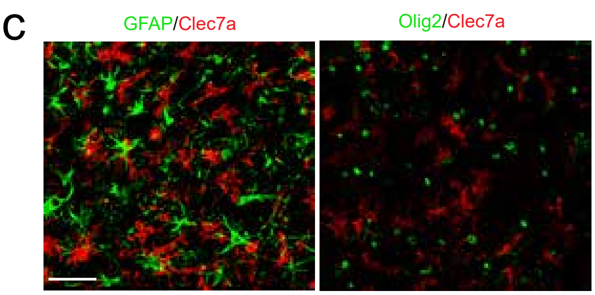


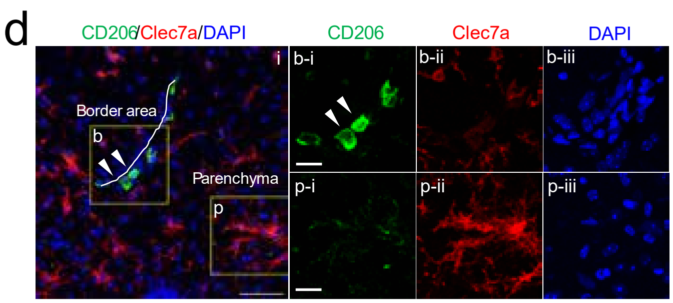


e

| Sample ID | Diagonosis | Age(y) | Gender | Clec7a in SC |
|---|---|---|---|---|
| 120 | FALS/FTD (C9) | 64 | M | + |
| 126 | FALS (C9) | 70 | M | + |
| 139 | FALS (C9) | 60 | M | + |
| 114 | FALS (SOD1) | 48 | M | + |
| 124 | FALS (SOD1) | 45 | F | + |
| 129 | FALS (SOD1) | 69 | M | - |
| 110 | SALS | 53 | M | + |
| 121 | SALS | 67 | M | + |
| 142 | SALS | 79 | M | - |
| 143 | SALS | 67 | M | - |
| 141 | SALS | 63 | M | - |
| 132# | sALS | 64 | F | - |
| 134# | sALS | 78 | F | + |
| 138# | sALS | 65 | M | + |
| 73# | Control | 74 | M | - |
| 83# | Control | 63 | F | - |
| 103# | Control | 92 | F | - |
| GBB-23-11 | Control | 62 | M | - |
| GBB-23-12 | Control | 67 | M | - |
| GBB-22-01 | Control | 47 | M | - |
| GBB-21-20 | Control | 89 | M | - |
| GBB-21-03 | Control | 71 | M | - |
| GBB-18-01 | Control | 68 | M | N/A |

Extended Data Fig. 1

**Extended Data Fig. 1 Selective up-regulation of Clec7a in microglia in ALS models and human ALS spinal cords**

**a.** Representative immunoblots of Clec7a and human SOD1 expression in the spinal cord tissues of WT, Clec7a KO, and SOD1G93A mice at pre-disease (P70), disease onset (P90), mid-disease (~P115), and end-disease (~P140) stages. **b.** Quantification of Clec7a expression normalized by β-actin in WT and SOD1G93A mice; n=3 mice/group. p value determined by one-way ANOVA and post-hoc Tukey's analysis. **c.** Representative images (40x) of Clec7a, GFAP, and Olig2 immunostaining in the spinal cord of SOD1G93A mice at mid-disease stage (P115). Scale bar: 50μm. **d.** Representative images (40x) of Clec7a and CD206 immunostaining in the spinal cord of SOD1G93A mice at mid-disease stage (P115); Scale bar: 50μm. bi-biii: magnified views of the box b in i; pi-piii: magnified views of the box p in i. box b and p represent border and parenchyma areas of the brain. Scale bar: 50μm (i) and 20μm (bi-biii, pi-piii). White arrows: $CD206^{+}$ border associate macrophages (BAMs) locating at ventral median fissure, indicated by the white line. **e.** Clinical information of human control and ALS cases from which spinal cords were from.

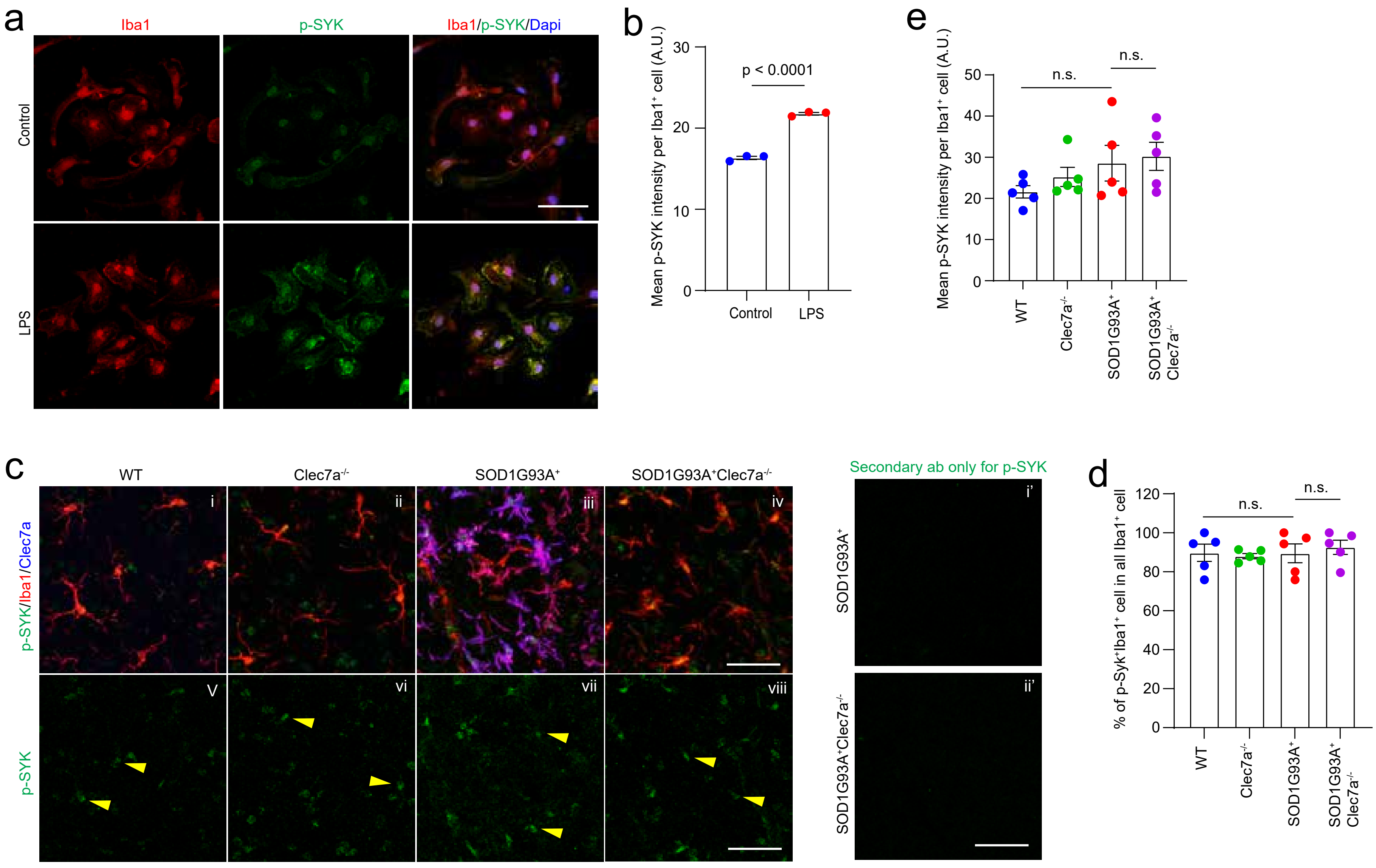


Extended Data Fig. 2

**Extended Data Fig. 2. Disease induces limited SYK activation in microglia that is not Clec7a dependent on spinal cords in SOD1G93A and SOD1G93AClec7a$^{-/-}$ mice**

Representative images (63x, **a**) of p-Y352 SYK and Iba1 immunostaining and quantification (**b**) of p-SYK intensity per Iba1$^{+}$ cells in LPS-treated primary microglia. n = 3 images/culture, 3 independent cultures. Scale bar: 50μm. Primary microglia cultures were treated with 10μg/mL LPS for 2hr. P value was determined by unpaired Student's t-test. **c.** Representative images (40x) of phosphorylated SYK (p-Y352 SYK), Iba1, Clec7a immunostaining in spinal cords of WT, Clec7a$^{-/-}$, SOD1G93A$^{+}$ and SOD1G93A$^{+}$Clec7a$^{-/-}$ mice at mid-disease stage (P115). i-iv: p-SYK, Iba1, and Clec7a immunoreactivity in all experimental groups. v-viii: p-SYK immunoreactivity in all experimental groups. i'-ii': secondary antibody (ab) only control for p-SYK immunostaining. Scale bar: 50μm. yellow arrows: p-SYK immunostaining in Iba1$^{+}$ cells. Quantification of the percentage (**d**) of p-SYK$^{+}$Iba1$^{+}$ in all Iba1$^{+}$ cells and mean p-SYK intensity (**e**) in Iba1$^{+}$ cells in experimental groups. n = 3 images/2-3 sections/mouse, 5 mice/group. P value was determined by one-way ANOVA and post-hoc Tukey's analysis. A.U.: arbitrary unit.

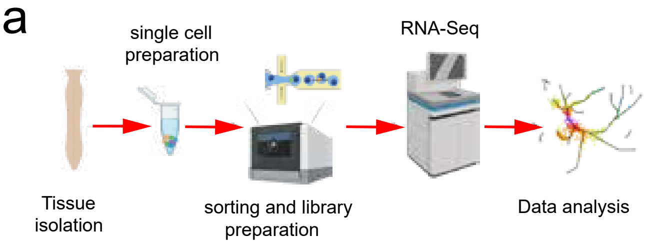


b

| Cell type | WT | Clec7a$^{-/-}$ | SOD1G93A$^{+}$ | SOD1G93A$^{+}$Clec7a$^{-/-}$ | Total numbers |
|---|---|---|---|---|---|
| Microglia | 6,745 | 10476 | 17,593 | 23,178 | 57,992 |
| Endothelia cell | 4169 | 5773 | 3588 | 3474 | 17,004 |
| Ependymal cell | 2745 | 1892 | 953 | 2126 | 7,716 |
| Astrocyte | 1229 | 1661 | 825 | 1062 | 4,777 |
| Oligodendrocyte | 978 | 1330 | 1210 | 1055 | 4,573 |
| Pericyte | 539 | 655 | 392 | 434 | 2,020 |
| Neuron | 312 | 433 | 234 | 205 | 1,184 |
| Fibroblast | 27 | 28 | 26 | 21 | 102 |
| Total | 16,744 | 22,248 | 24,821 | 31,555 | 95,368 |

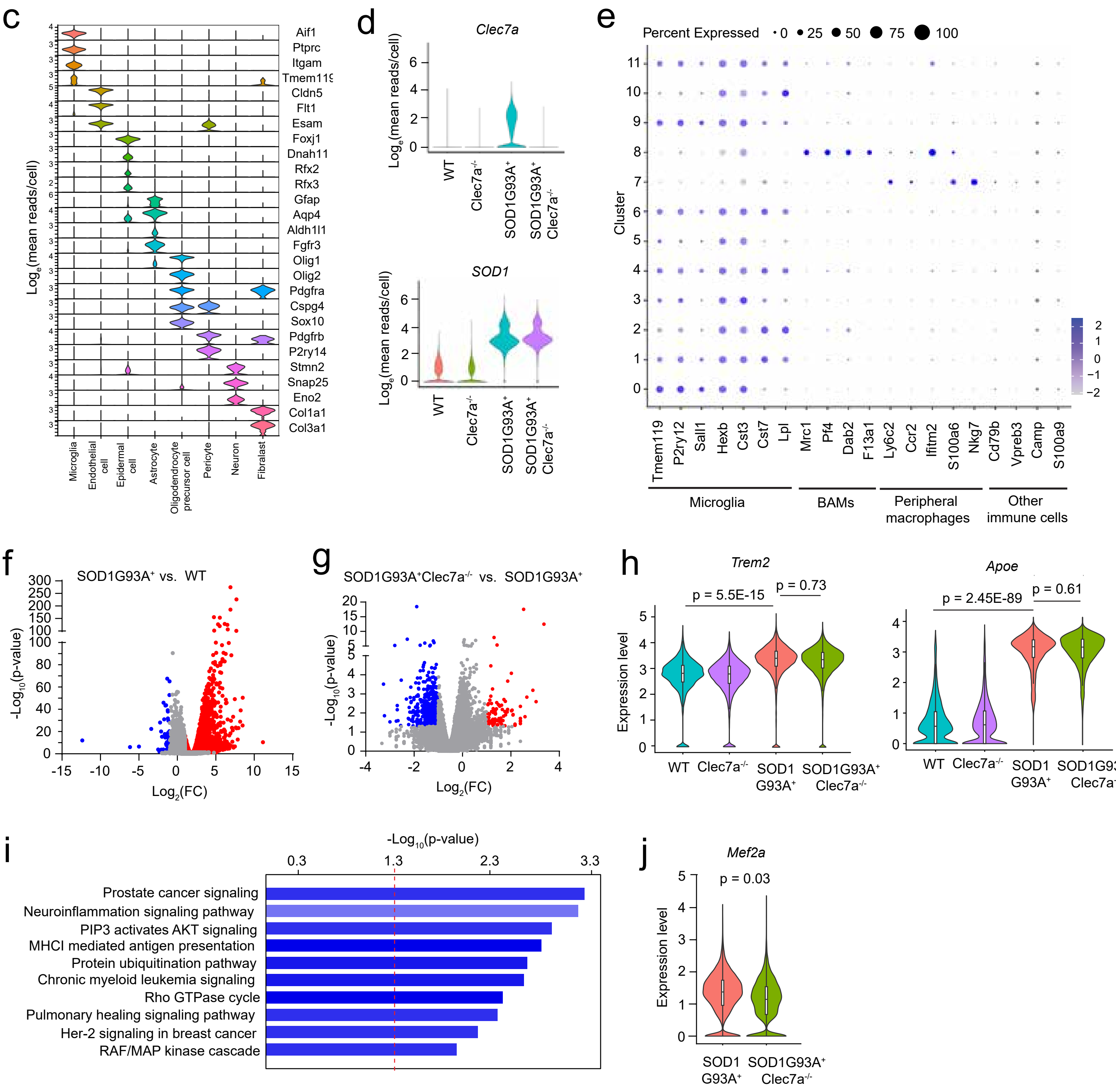


Extended Data Fig. 3

**Extended Data Fig. 3 Single cell RNA Sequencing (scRNA-seq) on spinal cords of SOD1G93A ALS mice**

**a.** Schematic diagram showing the workflow of performing scRNA-seq on spinal cord tissues from experimental mice. **b.** Total cell counts of each cell population identified in the spinal cord across four experimental groups from scRNA-seq. n = 3-4 mice/group. **c.** Stacked violin plot of cell signature gene expressions across different cell populations in spinal cords. **d.** Violin plots of *Clec7a* and *SOD1* expression among four experimental groups. **e.** Dot plot of lineage-specific marker expressions for microglia, BAMs, peripheral macrophages, and other immune cells, across 12 subclusters of microglia-like population. Volcano plot of microglial gene expression changes between WT and SOD1G93A mice (**f**) and between SOD1G93A$^{+}$ and SOD1G93A$^{+}$Clec7a$^{-/-}$ mice (**g**) based on pseudo bulk analysis. blue and red: significantly up- or down-regulated genes determined by the following criteria: $p < 0.05$, fold change (FC) > 2 or < -2, $\log_e$ (mean gene reads per sample) > 0.5. **h.** Violin plot of *Trem2* and *Apoe* expression in microglia in all experimental groups. **i.** Top 10 canonical signaling pathways enriched in DEGs found in SOD1G93A$^{+}$Clec7a$^{-/-}$ compared to SOD1G93A$^{+}$ microglia based on IPA pathway analysis. **j.** Violin plot of Mef2a expression in microglia in SOD1G93A$^{+}$ and SOD1G93A$^{+}$Clec7a$^{-/-}$ groups.

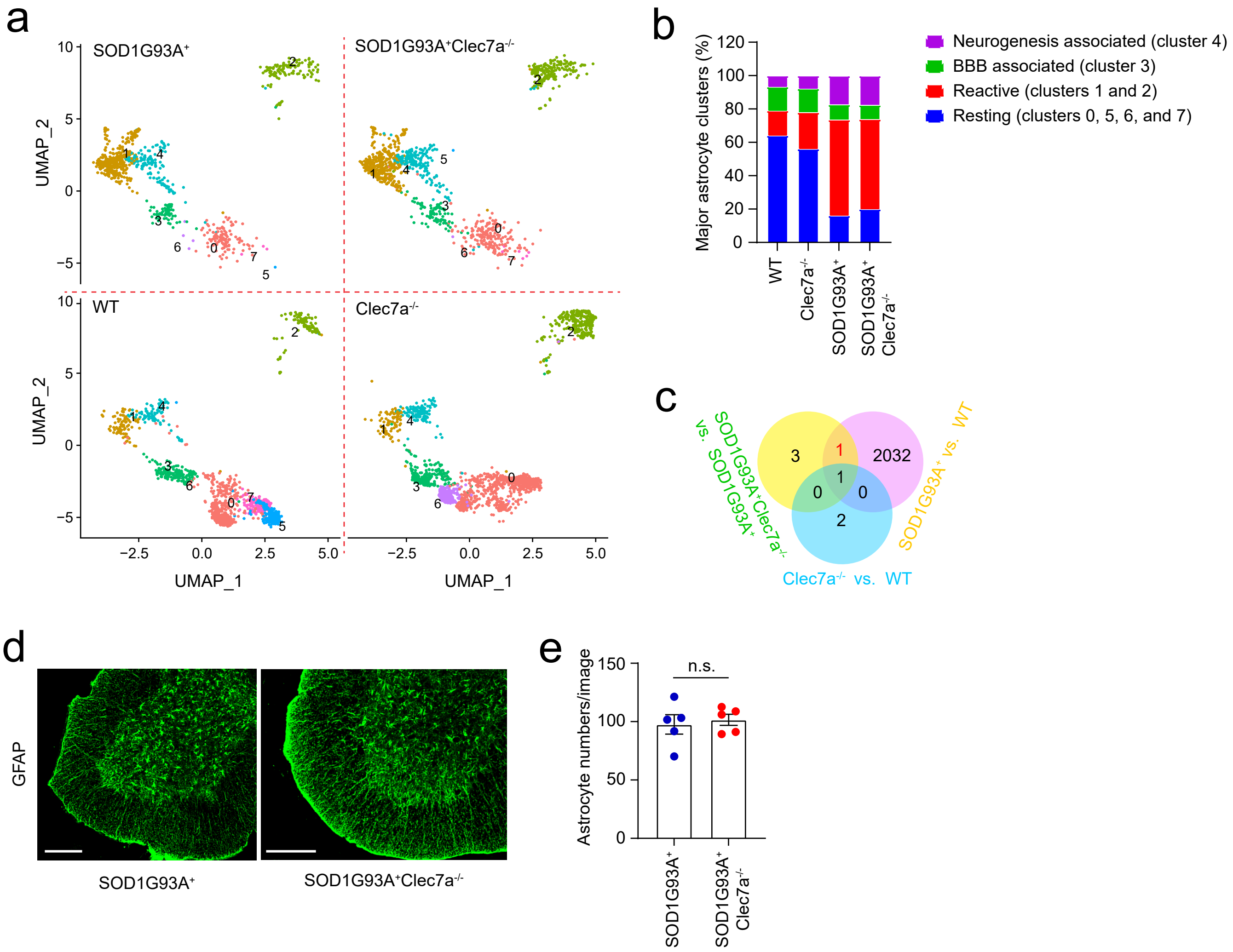


Extended Data Fig. 4

**Extended Data Fig. 4 Disease astrocyte gene expression changes are minimally affected by the deletion of CLec7a in SOD1G93A mice**

**a.** UMAP plots of astrocyte subclusters identified from scRNA-seq across four experimental groups. **b.** the percentage changes of major astrocyte subclusters from all experimental groups. **c.** Three-way Venn diagrams of differentially expressed genes (DEGs) in astrocytes between Clec7a$^{-/-}$ vs WT (blue), SOD1G93A$^{+}$ vs WT (pink), and SOD1G93A$^{+}$Clec7a$^{-/-}$ vs SOD1G93A$^{+}$ (yellow) from scRNA-seq. **d.** Representative images (20x) of GFAP immunostaining in the spinal cord sections of SOD1G93A$^{+}$ and SOD1G93A$^{+}$Clec7a$^{-/-}$ mice. Scale bar: 200μm. **e.** Quantification of GFAP$^{+}$ astrocyte numbers per image in spinal cords of SOD1G93A$^{+}$ and SOD1G93A$^{+}$Clec7a$^{-/-}$ mice. n= 6 images/3-4 sections/mouse, 5 mice/group. n.s.: not significant; p value determined by unpaired Student's t-test.

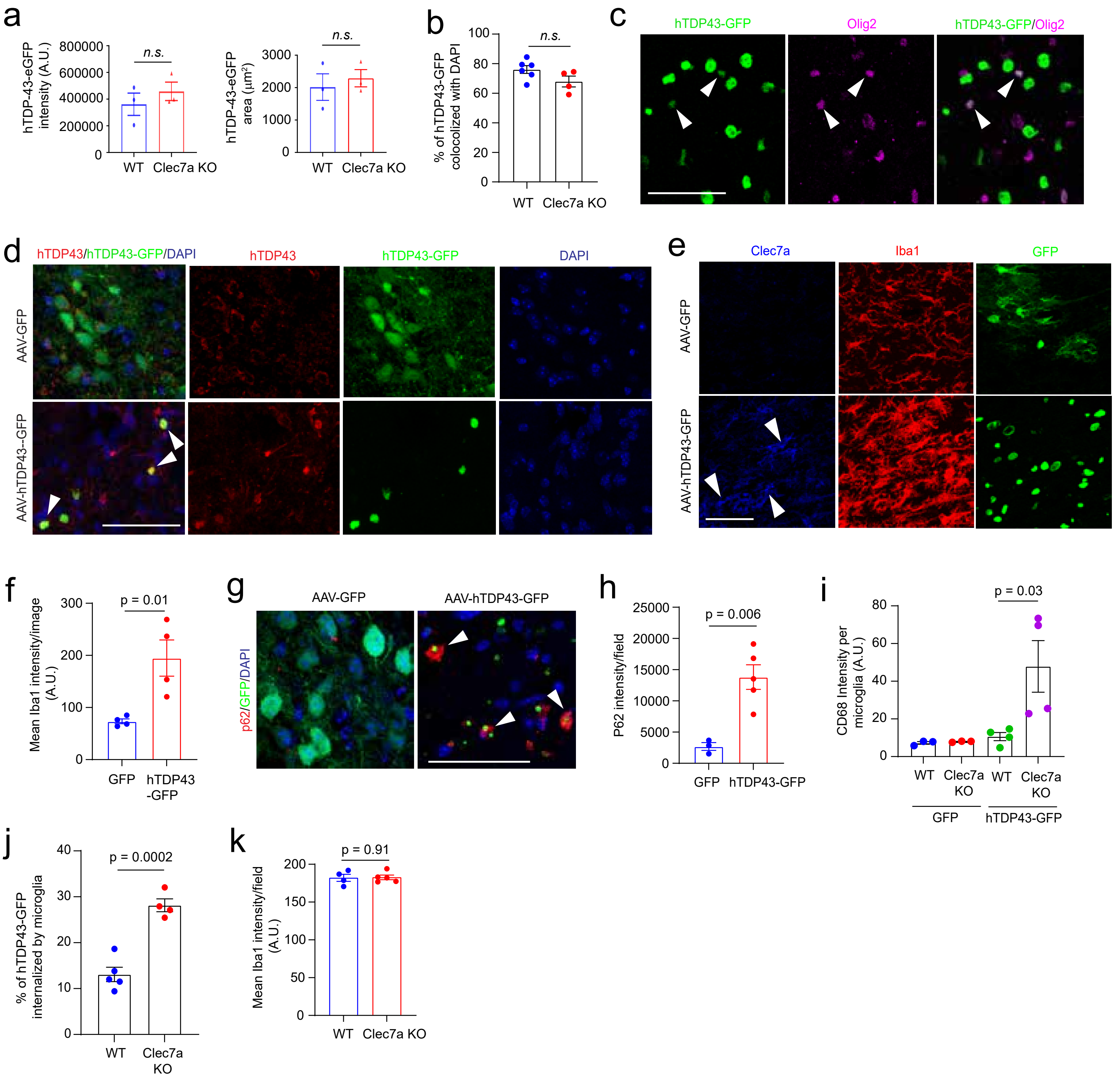


Extended Data Fig. 5

**Extended Data Fig. 5 Clec7a-deficient microglia promote phagocytosis of pathological hTDP43**

**a.** Quantification of hTDP43-GFP intensity and area per image following AAV9-hTDP43-GFP injection in the motor cortex of WT and Clec7a KO mice. n= 6-8 images/4-6 sections/mouse, 3 mice/group. n.s.: not significant; p value determined by unpaired Student's t-test. **b.** Quantification of the percentage of hTDP43-GFP$^+$ area colocalized with DAPI $^+$ area in AAV9-hTDP43-GFP-injected WT and Clec7a KO mice. n =6-8 images/4-6 sections/mouse, 4-6 mice/group. **c.** Representative images of Olig2 immunostaining on AAV9-hTDP43-GFP-injected mouse motor cortex. Scale bar: 50μm. **d.** Representative images of human TDP43 immunostaining in AAV-GFP and AAV-hTDP43-GFP-injected WT mouse motor cortex. White arrows: hTDP43 immunoreactivity colocalized with hTDP43-GFP puncta. Scale bar: 50μm. **e.** Representative images of Clec7a and Iba1 immunostaining in AAV-GFP and AAV-hTDP43-GFP-injected WT mouse motor cortex. Merged images are in Fig. 4b. Scale bar: 50μm. **f.** Quantification of Iba1 immunoreactivity per image from AAV-GFP and AAV-hTDP43-GFP-injected WT mouse motor cortex. n = 3-6 images/3 sections/mouse, 4 mice/group. Representative images of P62 immunostaining (**g**) and quantification of P62 immunoreactivity (**h**) in AAV-GFP and AAV-hTDP43-GFP-injected WT mouse motor cortex. white arrows: P62$^+$hTDP43-GFP$^+$ cells. n = 3-7 images/3-4 sections/mouse, 3-5 mice/group. p value determined by unpaired Student's t-test. **i.** CD68 intensity per microglia in the motor cortex of AAV-GFP and AAV-hTDP43-GFP-injected WT and Clec7a KO mice. n = 6-8 images/ 3-4 sections/mouse, 3-4 mice/group. p values were determined by one-way ANOVA and post-hoc Tukey's analysis. **j.** Quantification of the percentage of hTDP43-GFP puncta internalized by Iba1$^+$ microglia in WT and Clec7a KO. n = 6-12 images/4-6 sections/mouse,

4-5 mice/group. Scale bar: 50μm. p value determined by unpaired Student's t-test. **k.** Quantification of Iba1 immunoreactivity per image in the motor cortex of AAV-hTDP43-GFP-injected WT and Clec7a KO mice. n = 5-14 images/4-6 sections/mouse, 4-5 mice/group. p value determined by unpaired Student's t-test.

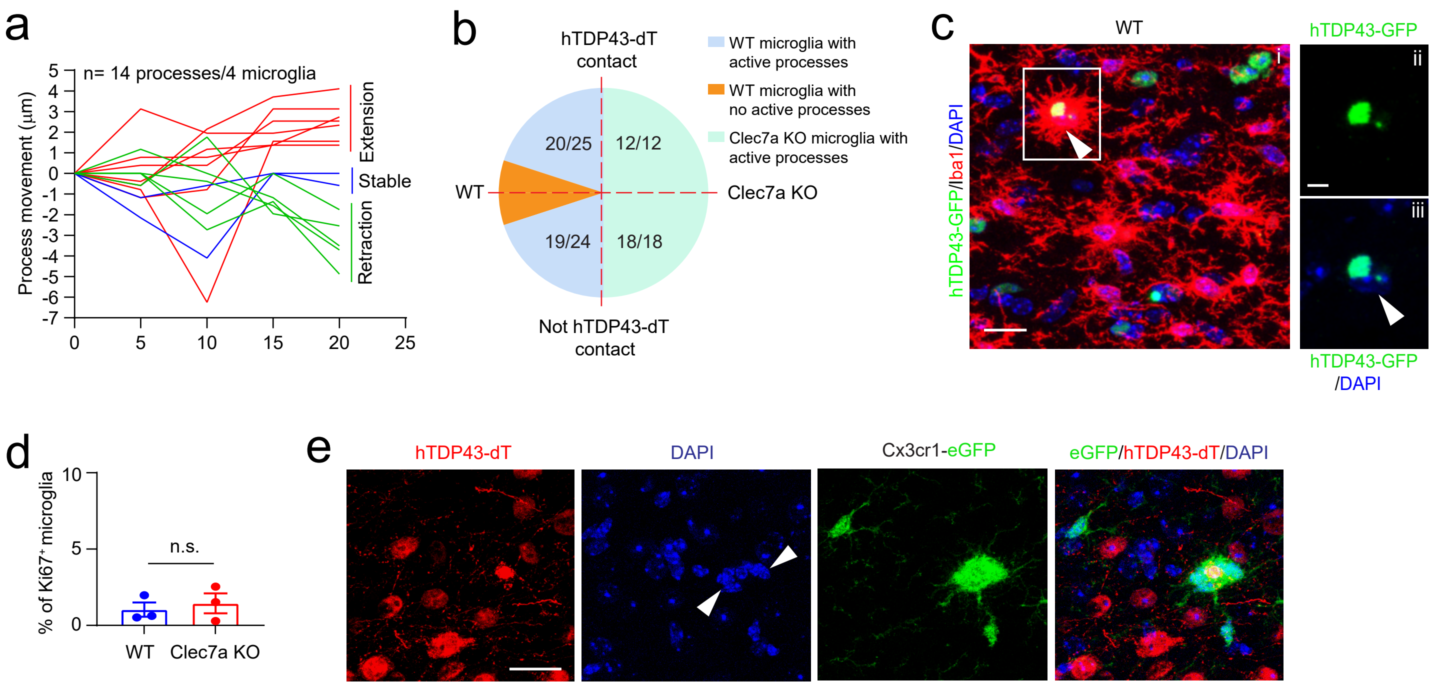


Extended Data Fig. 6

**Extended Data Fig. 6 Clec7a deficiency affects microglial process dynamics and microglial clustering around hTDP43**

**a.** Representative distribution of process movement during the 20-minute 2photon imaging session. Extension, stable, and retraction processes were determined by subtracting process length from 20 minutes (20’) to 0 minute (0’). Extension: 20’-0’ length > 1μm, stable: -1μm < 20’-0’ length < 1μm; retraction: 20’-0’ length < -1μm. A total of 14 processes from 4 microglia of one mouse were shown. **b.** Pie chart of the distribution of WT and Clec7a-deficient microglia with active process movement with or without contact with hTDP43-dT. n = 12-25 microglia/3-5 mice. **c.** Representative images showing hTDP43-GFP phagocytosis by single microglia in AAV-hTDP43-GFP-injected WT mouse motor cortex. Scale bar: 50μm. **d.** Quantification of the percentage of Ki67$^{+}$ microglia in the motor cortex of AAV-hTDP43-GFP-injected WT and Clec7a KO mice. n = 3-6 images/2-3 sections/mouse, 3 mice/group. p value determined using unpaired Student’s t-test. **e.** Representative images of multiple Cx3cr1-GFP$^{+}$ microglia surround hTDP43-dT for its phagocytosis and clearance in the motor cortex of AAV-hTDP43-dT-injected Clec7a KO mice. Scale bar: 20μm.

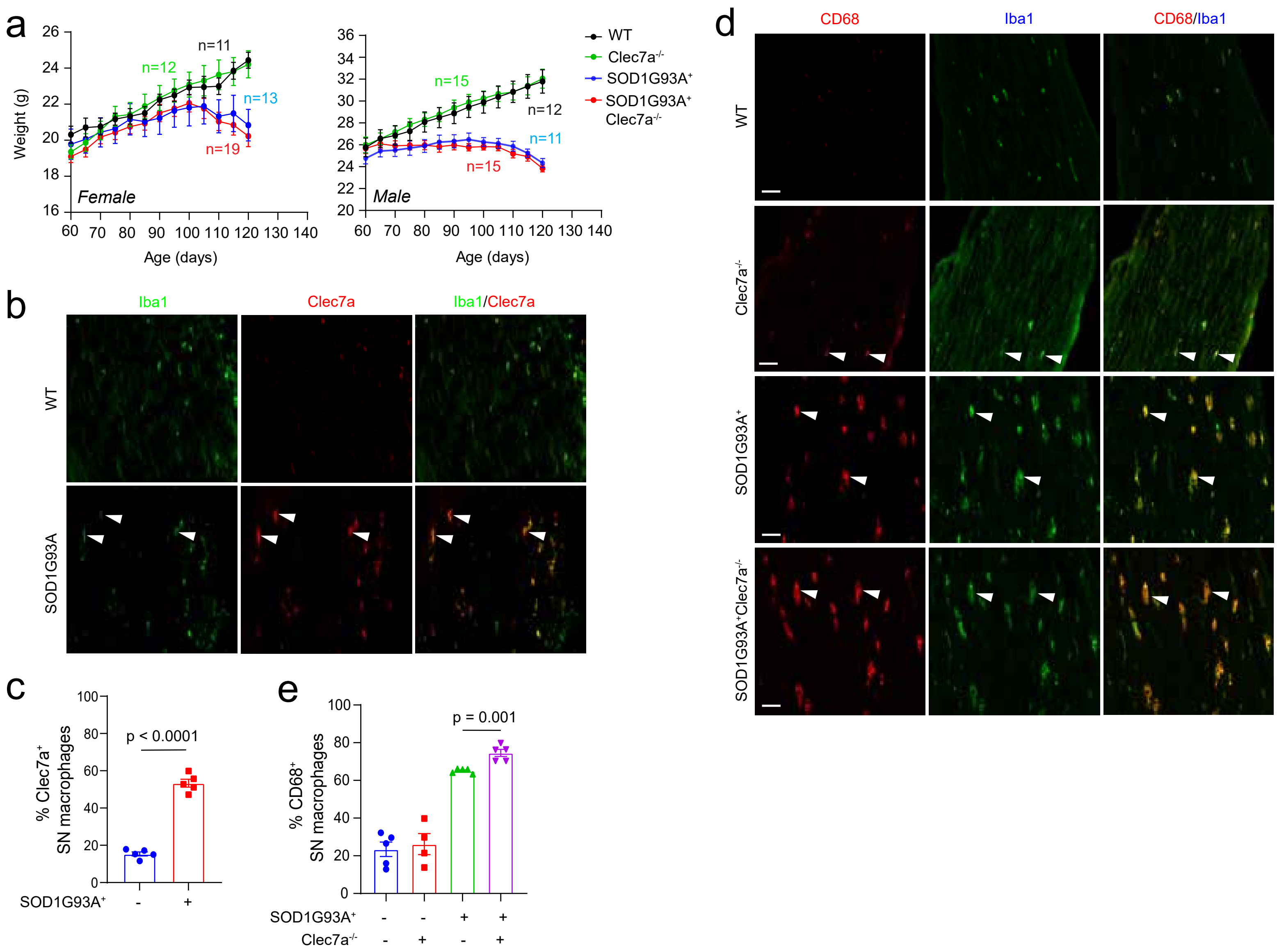


Extended Data Fig. 7

**Extended Data Fig. 7 Clec7a expression in sciatic nerve (SN) macrophages of SOD1G93A and SOD1G93A$^{+}$Clec7a$^{-/-}$ mice**

**a.** Weight growth plots of male and female mice from four experimental groups starting at 10 weeks. Number of mice in each group and sex were labeled in the same color as the group on the graph. Representative images (20x) of Clec7a immunostaining (**b**) and quantification of Clec7a immunoreactivity (**c**) in SN of WT and SOD1G93A mice at pre-symptomatic stage (P80). p value determined by unpaired Student's t-test. White arrows: Clec7a$^{+}$Iba1$^{+}$ macrophages. Scale bar: 50μm. n = 20-40 images/5-10 sections/mouse, 5 mice/group. **d.** Representative images (20x) of CD68 and Iba1 immunostaining (c) and quantification of number of CD68$^{+}$ macrophages (**e**) on SN of WT, Clec7a$^{-/-}$, SOD1G93A$^{+}$, and SOD1G93A$^{+}$Clec7a$^{-/-}$ mice at pre-symptomatic stage (P80). White arrows: CD68$^{+}$Iba1$^{+}$ macrophages. Scale bar: 50μm. n = 20-40 images/5-10 sections/mouse, 5 mice/group. p values determined by one-way ANOVA and post-hoc Tukey's analysis.

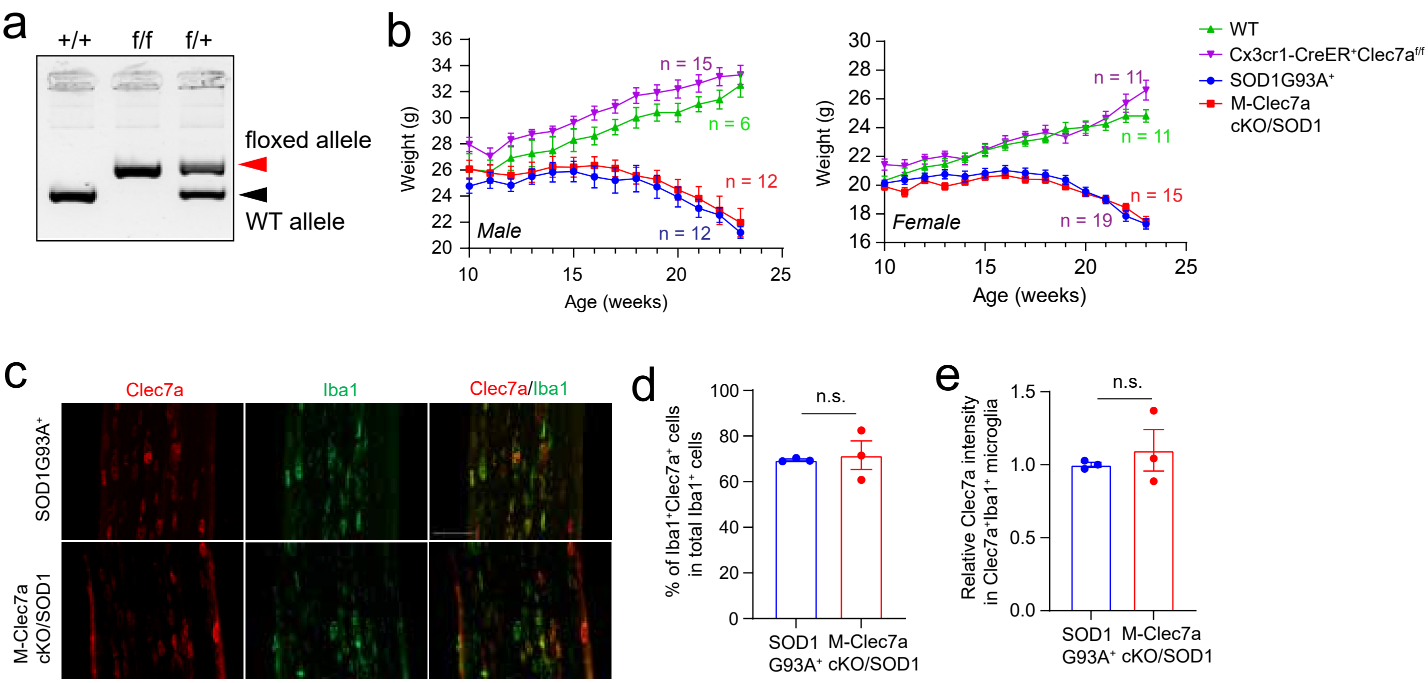


Extended Data Fig. 8

**Extended Data Fig. 8 Generation and validation of M-Clec7a cKO/SOD1 mice**

**a.** Representative genotyping PCR to identify Clec7af/f and f/+ alleles. Red and black arrows indicate floxed and WT allele respectively. **b.** Weight growth plots of male and female mice from four experimental groups starting at 10 weeks. Number of mice in each group and sex were labeled in the same color as the group on the graph. Representative Clec7 and Iba1 immunostaining images (20x, **c**) and quantification of Clec7a$^{+}$Iba1$^{+}$ SN macrophages numbers (**d**) and Clec7a immunoreactivity (**e**) in control SOD1G93A and M-Clec7a cKO/SOD1 mice. Scale bar: 100μm. n = 10-13 images/3-4 sections/mouse, 3 mice per group. p value determined by unpaired Student's t-test.

**Supplemental Table 1.** Differentially expressed genes in microglia between WT and SOD1G93A mice.

**Supplemental Table 2.** Differentially expressed genes in microglia between SOD1G93A and SOD1G93A Clec7a KO mice.

**Supplemental Table 3.** Clec7a regulated microglial differentially expressed genes (Clec7a-M-DEGs) that were up-regulated in SOD1G93A$^{+}$ relative to WT mice but were down-regulated in SOD1G93A$^{+}$Clec7a$^{-/-}$ relative to SOD1G93A$^{+}$ mice.

**Supplemental Table 4.** Differentially expressed genes in microglia between WT and Clec7a KO mice.

**Supplemental Table 5.** Gene promoters that bind to different transcription factors.

**Supplemental Table 6.** Differentially expressed genes in astrocytes between WT and SOD1G93A mice.

**Supplemental Table 7.** Differentially expressed genes in astrocytes between SOD1G93A and SOD1G93A Clec7aKO mice.

**Supplemental Table 8.** Differentially expressed genes in astrocytes between WT and Clec7aKO mice.
